# Salt-induced osmotic stress remodels osmoadaptive gene expression and physiology in the polyhydroxyalkanoate-accumulating thermophilic bacterium *Caldimonas thermodepolymerans*

**DOI:** 10.64898/2026.07.01.735907

**Authors:** Mohamed Mostafa, Radwa Moanis, Kristyna Hermankova, Yannick Gansemans, Rani Baes, Filip Van Nieuwerburgh, Karel Sedlar, Eveline Peeters

## Abstract

*Caldimonas thermodepolymerans* is a thermophilic polyhydroxyalkanoate (PHA)-producing bacterium with strong potential for sustainable bioplastic production. Besides serving as intracellular carbon and energy storage compounds, PHAs are increasingly associated with bacterial stress resistance and cellular robustness. This study aimed to investigate the physiological and transcriptomic response of *C. thermodepolymerans* to osmotic stress induced by elevated NaCl concentrations. Growth analysis demonstrated tolerance up to a supplementation of 2% NaCl, while moderate salt concentrations enhanced PHA accumulation, reaching 65% cell dry weight at 1.5% NaCl supplementation. To better understand the bacterial response to osmotic stress, RNA sequencing was performed under sublethal salt stress conditions. Differential expression analysis revealed major changes in genes related to osmoprotection, trehalose metabolism and type VI secretion systems, whereas motility and chemotaxis genes were strongly repressed. Phenotypic assays confirmed increased biofilm formation and reduced swarming motility under salt-induced osmotic stress. Although canonical PHA biosynthesis genes were not significantly differentially expressed, increased polymer accumulation suggests other underlying mechanisms linked to osmoadaptation. Together, these findings demonstrate that osmotic stress induces metabolic, physiological and regulatory responses in *C. thermodepolymerans*, highlighting the importance of PHA in stress adaptation besides its industrial applicability.

## 1. Introduction

The Gram-negative bacterium *Caldimonas thermodepolymerans* (formerly *Schlegelella thermodepolymerans*) is a moderately thermophilic organism with an optimum growth temperature between 50°C and 55°C (Elbanna et al. 2003; Kourilova et al. 2020). It has a high-GC genome (Musilova et al. 2021) and recently, efforts have been undertaken to develop a genome editing toolbox (Grybchuk-Ieremenko et al. 2025). *C. thermodepolymerans* is of interest to industrial biotechnology as it produces poly(3-hydroxybutyrate) (PHB), a polyhydroxyalkanoate (PHA) polyester, to over 80% of its cell dry weight (Kourilova et al. 2020; Zhou et al. 2023). PHAs stand out as a promising biodegradable alternative to petroleum-based plastics (Getino et al. 2024). In *C. thermodepolymerans*, PHA synthesis is performed via acyl-CoA dehydrogenase, enoyl-CoA hydratase, CoA transferase and a class I PHA synthase (Zhou et al. 2023). In addition to PHB synthesis, *C. thermodepolymerans* also secretes extracellular PHA depolymerase enzymes capable of PHA degradation (Elbanna et al. 2003; Kourilova et al. 2020).

*C. thermodepolymerans* exhibits a versatile carbohydrate catabolism, as its genome encodes Embden-Meyerhof-Parnas, pentose phosphate and Entner-Doudoroff pathways (Kourilova et al. 2020; Musilova et al. 2023). It preferentially utilizes D-xylose over D-glucose and is also capable of catabolizing L-arabinose, D-mannose, D-galactose and D-cellobiose thereby rendering the microbial species suitable for growth on lignocellulosic feedstocks (Zhou et al. 2025; Ozturk et al. 2025; Grybchuk-Ieremenko et al. 2025). *C. thermodepolymerans* is also capable of converting ferulic acid, a side product of lignocellulosic breakdown, into the valuable products vanillyl alcohol and vanillic acid (Hrabalová et al. 2024). These metabolic capabilities make *C. thermodepolymerans* an attractive industrial host for PHA production, together with its thermophilic lifestyle (Ozturk et al. 2025). Indeed, bioprocesses performed in high-temperature conditions eliminate the need for strict sterilization measures while cooling costs are also minimized given the cultivation at elevated temperature (Koller 2017). Recently, high-cell density cultivation in fed-batch fermentation of *C. thermodepolymerans* has been demonstrated with optimized PHB productivity and yield (Jang et al. 2025). Nevertheless, productivity is not yet sufficiently high and requires further engineering.

In the context of improving PHA productivities, salt and osmotic stress conditions constitute an important factor that can be used as an operational lever. In diverse bacterial and archaeal species, osmotic stress caused by high-salt conditions affects PHA accumulation and alters monomer composition (Guzmán et al. 2009; Shrivastav et al. 2010; Passanha et al. 2014; Cui et al. 2017; Godard et al. 2020). Mild salinity stress has been shown to improve yields. For example, in *Bacillus megaterium*, raising the NaCl concentration to approximately 7% resulted in an increase of PHB production from 6% to about 30% cell dry weight (CDW) (Godard et al. 2020). Moreover, in a bioreactor process with mixed microbial cultures, osmotic down- and upshifts were both found to boost PHA production capacity (Giovanella et al. 2025).

Salt stress response is believed to be linked to an osmoprotective role of PHA synthesis. This is exemplified by the model PHA producer *Cupriavidus necator*, which has been shown to exhibit reduced plasmolysis, less membrane damage and decreased cytoplasm leakage in conditions of high NaCl concentrations as compared to PHA-negative strains (Sedlacek et al. 2019). It is hypothesized that PHA granules help preserve cellular integrity during osmotic shifts, while the associated 3HB monomers can act as compatible solutes. Indeed, several species show an increased 3HB fraction under specific salinity conditions (Giovanella et al. 2025). From the viewpoint of process sustainability and downstream recovery, salt stress is also relevant for PHA production. The use of halophilic or halotolerant production hosts enables the reduction of freshwater use by employing seawater or saline side streams instead. Moreover, hypo-osmotic cell lysis enables fast and simple PHA recovery, reducing the need for harsh chemical or mechanical disruption methods (Bosma et al. 2013; Koller 2017), as commonly implemented with halophiles like *Haloferax mediterranei* (Cui et al. 2017). With this in mind, osmolysis was engineered in *C. necator* through improved osmolar stress resistance via adaptive laboratory evolution, thereby enabling water-induced product recovery (Adams et al. 2023).

*C. thermodepolymerans* belongs to the order Burkholderiales and shares phylogenetic affinity with other well-characterized PHA-producing bacteria, including *C. necator* and *Burkholderia* species. Osmotic stress responses have been studied in other Burkholderiales members. *Burkholderia cenocepacia* employs diverse osmoadaptation strategies, of which the constitutive or inducible intracellular accumulation of compatible solutes, including glycine betaine, trehalose and amino acids, is a major mechanism (Behrends et al. 2011). In *C. necator*, the large-conductance mechanosensitive channel MscL has been shown to be involved in osmotic stress (Adams et al. 2023).

In contrast to other *Burkholderiales* and despite NaCl-induced stress being a key process parameter for PHA production, the impact of osmolarity stress on *C. thermodepolymerans* has remained uncharacterized thus far. In this study, we aim to bridge this knowledge gap by studying the physiological and genetic response of *C. thermodepolymerans* to increased NaCl concentrations. This is followed by a transcriptome analysis using RNA-sequencing to unravel which metabolic pathways and physiological processes are transcriptionally responsive to an elevated supplemented NaCl concentration of 2%. By revealing the NaCl-induced osmotic stress response of *C. thermodepolymerans* in relation to its PHA production capability and identifying key genetic factors in this stress response, we contribute to the development of novel strategies for improving its future use as an industrial PHA production host.

## 2. Materials and methods

### 2.1. Bacterial strain and growth conditions

The strain *C. thermodepolymerans* DSM 15344^T^ was obtained from the German Collection of Microorganisms and Cell Cultures (DSMZ) (Leibniz Institute, Braunschweig, Germany).

Liquid cultures were prepared in tryptic soy broth (TSB) medium (Sigma-Aldrich), which is composed of 17 g/L casein peptone (pancreatic), 2.5 g/L K_2_HPO^4^, 2.5 g/L glucose, 5 g/L NaCl and 3 g/L soya peptone (papain digest) (Smith and Dell 1990). Additional NaCl was added to TSB medium, leading to a range of supplemented NaCl concentrations from 0 to 2% (w/v), corresponding to total NaCl concentrations between 0.5% (w/v, no additional NaCl added) to 2.5% (w/v). Cultivation was performed at 55°C with shaking at 180 revolutions per minute (rpm) and growth was monitored by measuring the optical density at 600 nm (OD₆₀₀).

For cultivation on solid media, tryptic soy agar (TSA) was prepared by supplementing TSB with 15 g/L agar. Plates were incubated at 55°C.

For growth experiments, 7.5 mL of TSB medium with indicated supplemented NaCl concentration was prepared in 50 mL TubeSpin® Bioreactor tubes. These tubes were inoculated with precultures and the initial OD_600_ was set to 0.25-0.30. Subsequently, they were placed in the RTS-8 personal bioreactor system at 55°C, 1000 rpm with OD₆₀₀-equivalent measurements recorded every 3 minutes. These experiments were performed in biological triplicate for each condition.

For growth-curve modelling, three replicates per condition were fitted using the Baranyi primary growth model (Baranyi and Roberts 1994) implemented in R (version 4.3.1) using the nlsLM function from the minpack.lm package. Model fitting was performed by nonlinear regression for the exponential growth phase, determined by plotting lnOD_600_ versus time and the following parameters were estimated: lower asymptote (y₀), maximum population density (y_max_) and maximum specific growth rate (μ). Lag phase duration was estimated from the experimental data prior to modelling. Model performance and goodness of fit were evaluated using the root mean square error (RMSE) between observed and predicted OD_600_ values.

### 2.2. RNA extraction and RNA-sequencing

Cultures were pelleted and stabilized using RNAprotect Cell Reagent (Qiagen) and total RNA was extracted from the cell pellets using a Promega SV Total RNA Isolation System according to the manufacturer’s instructions with minor modifications. An additional lysis step was included using a freshly prepared Tris-EDTA buffer (10 mM Tris base and 1 mM EDTA-Na₂·2H₂O, pH 7.5) containing lysozyme. To ensure removal of genomic DNA, DNase treatment was performed twice: first by on-column DNase treatment according to manufacturer’s instructions, followed by an additional treatment using the TURBO DNA-free kit (Thermo Fisher Scientific). The total RNA quantity was determined with a Quant-iT RiboGreen RNA assay (Thermo Fisher Scientific) and RNA integrity was evaluated on a Bioanalyzer instrument with an RNA 6000 Nano chip (Agilent Technologies). Ribosomal RNA depletion was established using a PAN-Bacteria riboPOOL kit (siTOOLS), followed by purification with a Zymo RNA Clean and Concentrator-5 kit (Zymo Research). Four biological replicates were included for each condition: a control condition, consisting of TSB without supplemented NaCl, and the treatment condition, consisting of TSB medium with supplemented NaCl concentration of 2.0% (w/v).

Sequencing libraries were prepared with the xGen Reverse Stranded RNA Library Prep Kit for Illumina (Integrated DNA Technologies) in combination with the xGen UDI 8 nucleotide (nt) primer pairs for Illumina (Integrated DNA Technologies). A library enrichment PCR was performed for 14 cycles and the libraries were purified with 0.8x AMPure XP beads (Beckman Coulter). Additional purification of the libraries was performed on an Egel EX 2% agarose gel (Thermo Fisher Scientific) to remove residual adapter dimers. Library quality was verified on a Bioanalyzer instrument with a DNA High Sensitivity chip (Agilent Technologies) and concentrations were measured with qPCR according to the ‘Sequencing library quantification guide’ (Illumina). Sequencing was performed on a NextSeq500 instrument (Illumina) in high-output mode with 75 nt single-end reads and a 3% PhiX spike-in.

### 2.3. RNA-sequencing data analysis

Raw RNA-sequencing (RNA-seq) reads were trimmed to remove adapter sequences and low-quality bases using fastp (v1.0.1) (Chen 2025), with adapter sequences checked against TruSeq3-SE.fa from Trimmomatic (v0.39) (Bolger et al. 2014). Subsequently, additional *in silico* removal of rRNA transcripts was performed using SortMeRNA (v4.3.7) (Kopylova et al. 2012). The rRNA-depleted reads were then mapped to the genome of *C. thermodepolymerans* DSM 15344^T^ (Musilova et al. 2021) (annotation CP064338) using STAR (v2.7.11b) (Dobin et al. 2013). Each step of the RNA-seq data preprocessing, including initial quality assessment of raw reads, was inspected using FastQC (v0.12.1) (Andrews 2010) and summarized with MultiQC (v1.32) (Ewels et al. 2016).

Gene-level read counts were generated from mapped reads using featureCounts (Liao et al. 2014) from the Subread package (v2.0.6). Genes encoding rRNAs and tRNAs were excluded from downstream analyses. Differential expression analysis was performed using DESeq2 (v1.42.1) (Love et al. 2014). Statistical significance was assessed using an adjusted p-value (padj) threshold of 0.05, applying the Benjamini–Hochberg (BH) method for multiple testing correction. Genes were considered differentially expressed if they met the criteria of padj < 0.05 and |LFC| > 1 with LFC = log_2_ fold change. Functional enrichment analysis and small RNA (sRNA) analysis are described in **Supplementary Note S1**. Heatmaps were generated using the ComplexHeatmap (v2.18.0) (Gu 2022), while all other plots were created using the R package ggplot2 (v4.0.2) (Wickham 2016).

### 2.4. Polyhydroxyalkanoate extraction

PHA extraction was performed in five biological replicates as described previously (Mozejko-Ciesielska et al. 2017). A 100-mL aliquot of the bacterial culture was centrifuged at 10,000 x *g* for 10 minutes and the pelleted biomass was freeze-dried by submerging the sample in liquid nitrogen followed by lyophilization for 24 hours. Next, the sample was weighed for determination of the cell dry weight (CDW). PHA was extracted by incubating the dry lyophilized biomass in chloroform at 50°C for 3 hours while shaking at 180 rpm. The mixture was then filtered through No. 1 Whatman filter paper using a vacuum pump (Welch™ Standard Duty WOB-L™ Piston Vacuum Pump: Model 2522) and left to dry at room temperature, after which the formed PHA film was weighed.

The PHA content was calculated as the ratio of extracted PHA weight to CDW as follows (Johnston et al. 2018):

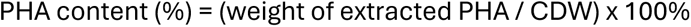

Statistical analysis was performed using one-way analysis of variance (ANOVA) followed by Tukey’s HSD test to evaluate the effect of different NaCl concentrations on PHA content. Differences were considered statistically significant at p < 0.05.

### 2.5. Mass spectrometry analysis of isolated polyhydroxyalkanoate granules

Cells were harvested from a 300-mL culture cultivated for 24 hours and intracellular PHA granules were purified by glycerol-gradient ultracentrifugation (Li et al. 2019). The visible PHA-containing layer formed between the glycerol phases was collected and subjected to an additional ultracentrifugation step to obtain a purified PHA granule pellet. The presence of PHA in the purified granule fraction was confirmed by Fourier-transform infrared spectroscopy (FTIR) using commercial PHBV as a reference (**Supplementary Note S2**).

Granule-associated proteins were analyzed using sodium dodecyl sulfate-polyacrylamide gel electrophoresis (SDS-PAGE) and shotgun liquid chromatography–tandem mass spectrometry (LC-MS/MS) analysis performed by the VIB Proteomics Core, Ghent University.

Mass spectrometry data were processed using MaxQuant software version 2.1.3.0 (Cox and Mann 2008). Protein identification was performed against the *C. thermodepolymerans* protein sequence database and protein abundance was estimated using intensity-based absolute quantification (iBAQ) values (Schwanhäusser et al. 2011). Proteins with the highest iBAQ values were considered the most abundant proteins associated with the granule preparation.

### 2.6. Motility assay

Solid motility-assay media were prepared by supplementing the corresponding liquid media with 0.7% agar and pouring into Petri dishes (Palma et al. 2022). Overnight-cultivated precultures were used to inoculate the plates in the centre, after which plates were incubated at 50°C for 24 hours. All assays were performed in biological triplicate. The formation of spreading growth after 48 hours of incubation was considered evidence for motility, compared to aggregation of growth at the site of inoculation. The diameter of the growth was recorded by measuring the widest distance across the growth zone passing through the centre.

### 2.7. Biofilm formation assay

Biofilm formation was quantified using the microtiter dish crystal violet assay, as described by (O’Toole 2011). *C. thermodepolymerans* precultures were diluted in fresh medium (1:100), of which 100 µL was transferred to a 96-well plate, which was incubated at 55°C during 7 hours. Each condition was studied in five replicates. Cultures were discarded and wells were washed twice with water. Subsequently, 125 µL of 0.1% crystal violet was added to each well followed by an incubation at room temperature for 15 minutes. The plate was washed again with water three to four times and dried overnight. To quantify biofilm formation, 125 µL of 30% acetic acid was added to each well and the plate was incubated again at room temperature for 15 minutes. A 125-μL aliquot of solubilized crystal violet was transferred to a new plate and absorbance was quantified at 550 nm (A_550_) using a BioTek Synergy H1 Microplate reader. Statistical analysis was performed using one-way analysis of variance (ANOVA) followed by Tukey’s HSD test with significance considered at p < 0.05.

## 3. Results

### 3.1. Establishing the salt stress window for Caldimonas thermodepolymerans

We first set out to assess how increasing NaCl concentrations affect growth physiology upon cultivating *C. thermodepolymerans* cells in TSB medium at 55°C (**Figure 1**; **Supplementary Table S1**). *C. thermodepolymerans* tolerated supplemented NaCl concentrations up to 2% (w/v) in TSB medium, although growth was progressively impaired at high salinities. Growth trajectories were comparable at supplemented NaCl concentrations of ≤1% (w/v), whereas 1.5% reduced both growth rate and final biomass and 2% strongly inhibited bacterial proliferation, consistent with a sublethal stress condition (**Figure 1**). Across conditions, three distinct growth phases were evident: a lag phase (µ < 0.1 h⁻¹), an exponential phase reaching a maximum specific growth rate (µ_max_) and a stationary phase beginning thereafter (µ < 0.09 h⁻¹). Baranyi model fits confirmed that an increasing supplemented NaCl concentration resulted in a decreased µ_max_ and was accompanied by longer lag phases and lower growth rates at ≥ 1.0% supplemented NaCl (**Supplementary Table S1**).

**Figure 1.**
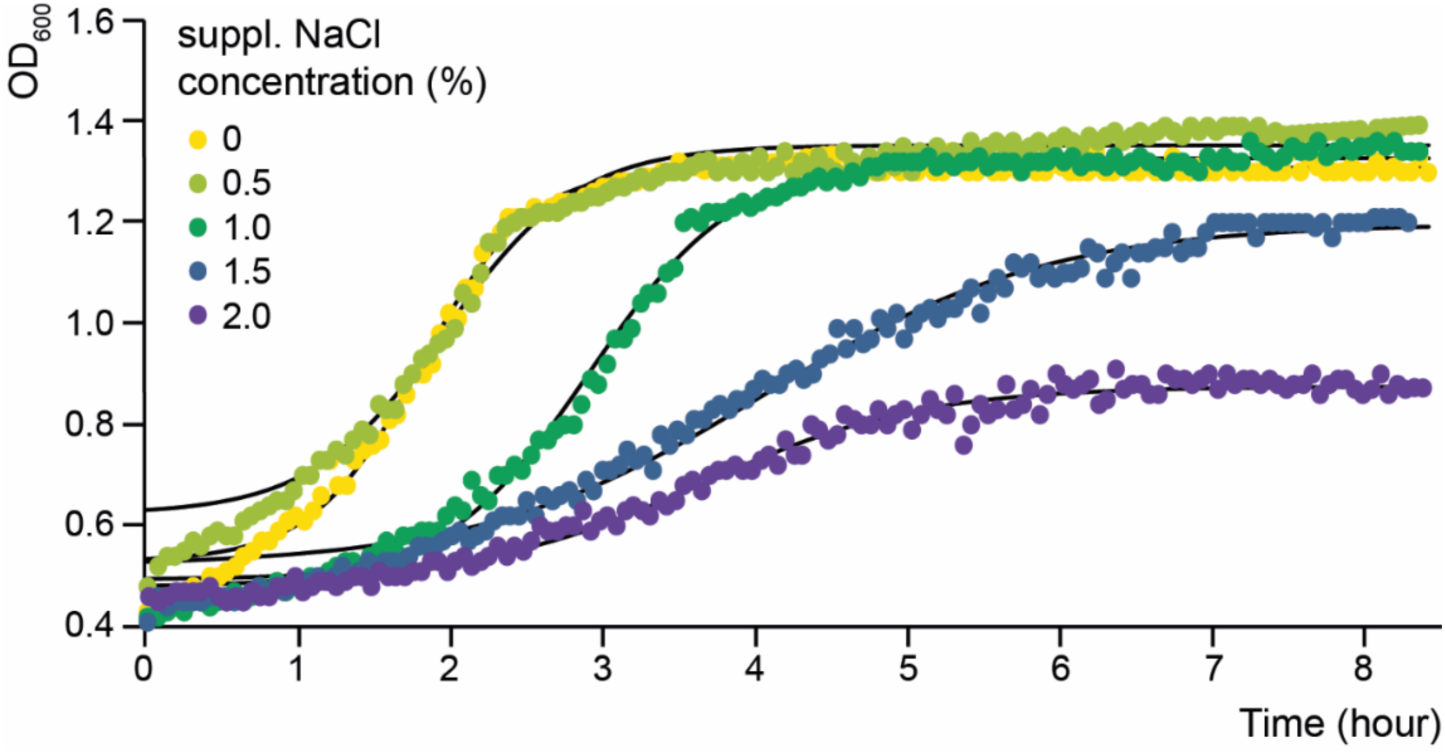
Growth kinetics of *C. thermodepolymerans* in TSB medium with 0–2% supplemented NaCl (w/v). Cultures were incubated at 55°C and growth was monitored as OD₆₀₀ over time. OD₆₀₀ values from one representative biological replicate are shown; experiments were performed in triplicate. Colored data points show measured OD₆₀₀ values and black lines show Baranyi model fits. Estimated growth parameters are shown in **Supplementary Table S1**.

### 3.2. Global transcriptomic response to 2% supplemented NaCl

As a sublethal but strong osmotic stress condition, the condition with 2% supplemented NaCl was selected for comparative RNA-seq analysis to identify genetic responses of *C. thermodepolymerans* to elevated salinity as compared to a control condition without supplemented NaCl. Cells were harvested for RNA sampling at mid-exponential phase and subsequent RNA-seq generated 21.8-25.8 million single-end reads per sample (**Supplementary Table S2**). After quality filtering, reads showed high quality (mean Phred score > 30) with lengths of 66-76 bp (**Supplementary Figure S1**) and a Principal Component Analysis (PCA) separated the NaCl stress and control conditions (**Supplementary Figure S2**). This transcriptomic profiling resulted in the identification of 764 differentially expressed genes (21.28% of total CDS) (padj < 0.05 and |LFC| > 1) out of 3,589 coding sequences (CDS) (**Supplementary Figure S3**; **Supplementary File S1**). Among these, 279 genes were downregulated (7.77% of total CDS) and 485 were upregulated (13.51% of total CDS) in response to increased salinity. This global response indicates substantial physiological remodelling under salt stress.

A Clusters of Orthologous Groups (COG) enrichment analysis was performed to assess the stress response on a broader functional level (**Figure 2**; **Supplementary File S2**). COG annotation assigned more than 70% of all CDS to functional categories with defined biological roles (**Supplementary Table S3**). Overrepresented functional classes (*i.e.* containing more differentially expressed genes than expected) were evaluated separately for up- and downregulated genes. This enrichment analysis identified significantly overrepresented terms across all three Gene Ontology (GO) domains: 16 terms related to biological processes (**Supplementary Figure S4**), 12 terms describing molecular functions (**Supplementary Figure S5**) and 8 terms associated with cellular components (**Supplementary Figure S6**). Functional categories enriched among upregulated genes included genes classified in the ‘Carbohydrate transport and metabolism’ category (G), ‘Cell wall/membrane/envelope biogenesis’ category (M) and the ‘Function unknown’ category (S), indicating that osmotic stress leads to adaptive responses with known functions, as well as a response involving poorly characterized genes. Conversely, only one category was enriched among downregulated genes, namely ‘Cell motility’ (N), suggesting a reduced investment in motility under salt stress. Several COG classes were underrepresented, indicating fewer differentially expressed than statistically expected in these categories, including ‘Energy production and conversion’ (C), ‘Translation, ribosomal structure and biogenesis’ (J), ‘Transcription’ (K) and ‘Replication, recombination and repair’ (L) (**Figure 2**). Together, this suggests that core housekeeping functions were comparatively less transcriptionally responsive to salt stress, even though growth was clearly impacted under high NaCl conditions (**Figure 1**).

**Figure 2.**
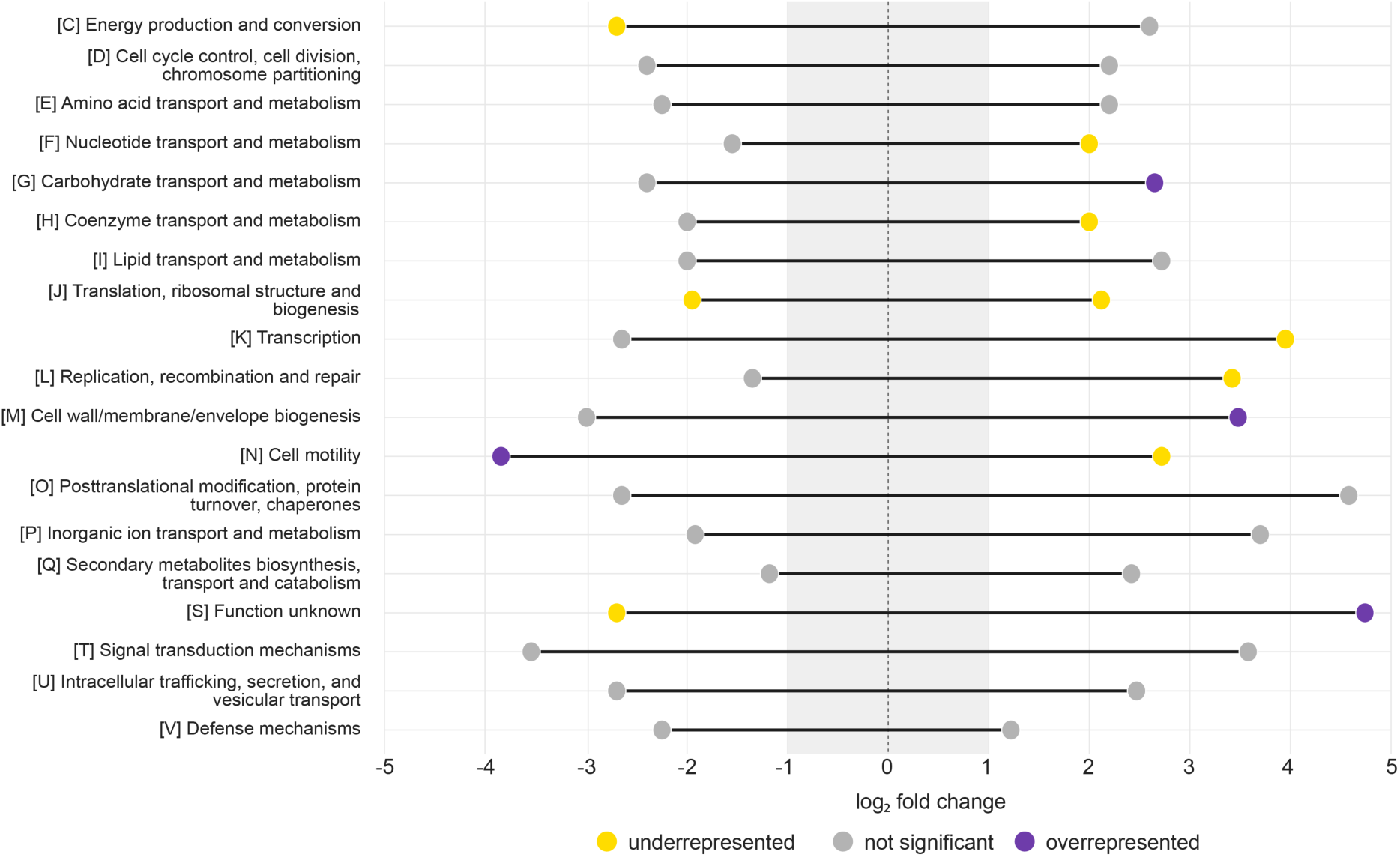
COG enrichment analysis visualized using a dumbbell plot. For each COG category, the range of log₂ fold change (LFC) values is shown, with minimum and maximum values indicated by the endpoints of each dumbbell. Differential expression analysis results were used to test whether each COG category was enriched among upregulated or downregulated genes. Overrepresented categories are shown in purple, underrepresented categories in yellow, and non-enriched categories in grey. The grey-shaded region from −1 to 1 denotes the applied filtering threshold (|log2 fold change| > 1). Points falling outside this region indicate genes that met the fold-change cutoff used to define differential expression.

Not only protein-encoding CDS sequences were differentially expressed under osmotic stress, but non-coding RNAs also showed marked differential expression, indicating an additional layer of post-transcriptional regulation. A total of 2,106 small RNAs (sRNAs) were predicted with a length between 75 and 1500 nt (Barik and Das 2018) with the majority falling within the range 75–700 nt (**Supplementary Figure S7**; **Supplementary File S3**). After filtering and classification, 2,060 *cis*-encoded and 46 *trans*-encoded sRNAs were predicted. Among these, 292 sRNAs were differentially expressed, dominated by *cis*-encoded sRNAs (285 differentially expressed, of which 220 and 65 up- and downregulated, respectively) with a smaller contribution of *trans*-encoded sRNAs (7 differentially expressed, of which 4 and 3 up- and downregulated, respectively). Representative examples of *cis*- and *trans*-encoded sRNAs, including their genomic organization and strand-specific RNA-seq expression profiles, are shown in **Supplementary Figures S8** and **S9**, respectively.

This filtering step resulted in 299 sRNA-target pairs that were further prioritized, and correlations between their transcriptional profiles were subsequently assessed. Operon-aware target assignment resulted in the prediction of 299 prioritized sRNA-target pairs, of which 75 showed statistically significant expression correlations (FDR<0.05 & |r|>0.5) (**Supplementary File S3**). This bias towards *cis*-encoded sRNAs suggests that antisense-linked regulation may be prevalent in *C. thermodepolymerans*.

### 3.3. Salt-induced osmotic stress uncouples polyhydroxyalkanoate accumulation from canonical *pha* gene expression

To assess the impact of salt-induced osmotic stress on PHA production, *C. thermodepolymerans* was cultivated in TSB medium with supplemented 0.5–2% (w/v) NaCl at 55°C for 48 hours, after which PHA was extracted and quantified (**Figure 3A**). PHA content increased with NaCl supplementation, reaching 65% CDW at 1.5% (w/v) NaCl as compared to 42.5% CDW for unsupplemented TSB medium. However, at 2% (w/v) NaCl, PHA accumulation declined sharply to 12.8% CDW, consistent with this concentration exerting an osmotic stress that impairs growth (**Figure 1**). This confirms that a supplementation of 2% (w/v) NaCl approaches the upper physiological tolerance limit of *C. thermodepolymerans.* Statistical analysis showed that PHA content at 1.5% NaCl was significantly higher than at 0.0%, 0.5%, and 1.0% NaCl, whereas 2.0% NaCl resulted in significantly lower PHA content than all other treatments.

**Figure 3.**
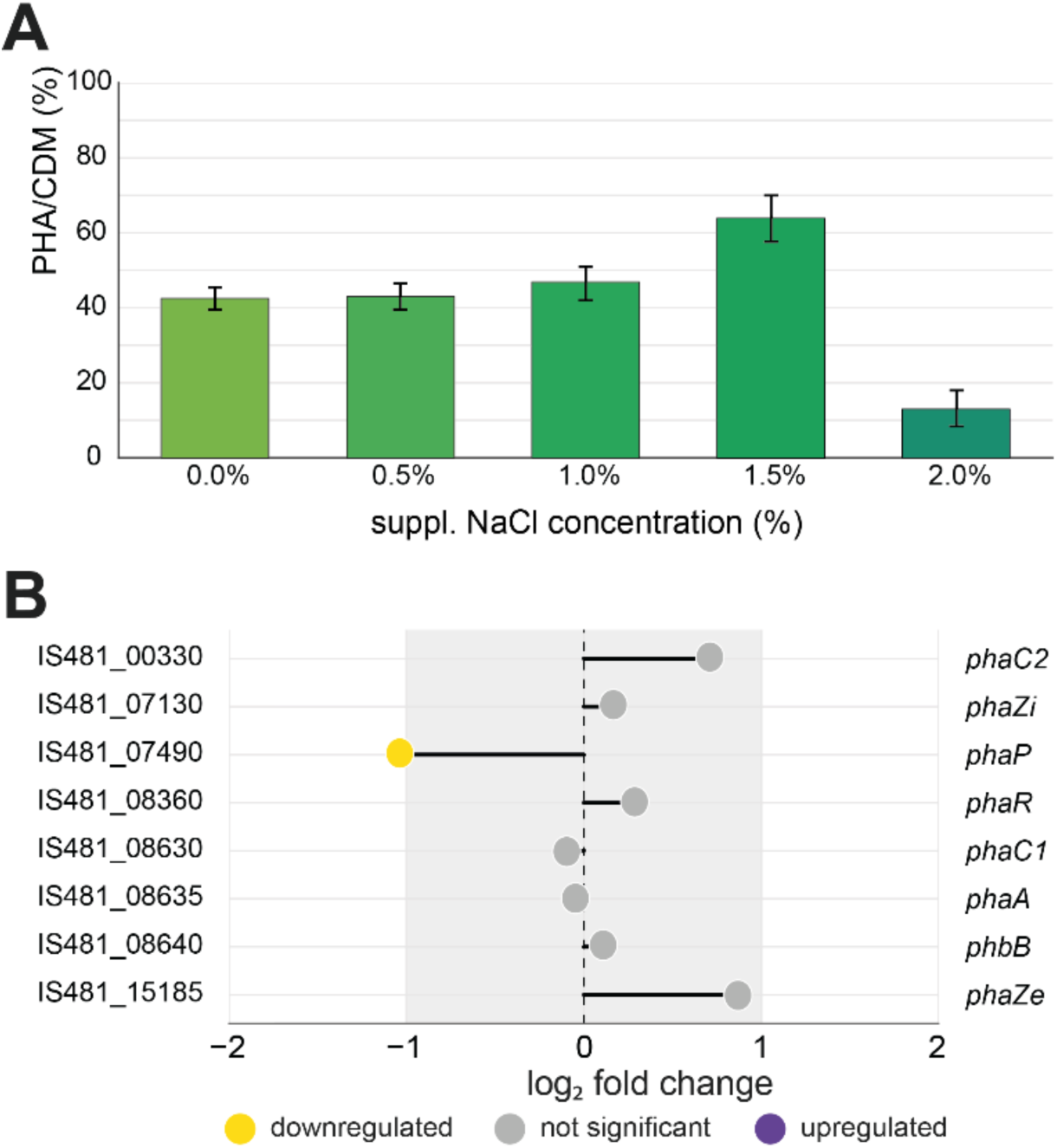
PHA accumulation and transcriptional response of PHA-associated genes in *C. thermodepolymerans* under NaCl stress. (**A**) PHA content was quantified as percentage of cell dry weight (% CDW) after cultivation in TSB with 0.0–2.0% (w/v) supplemented NaCl for 48 hours. Data represents biological replicates (n = 5), error bars show the standard deviation; Statistical significance was determined by one-way ANOVA with Tukey’s HSD test (*p* = 6.95E-12). (**B**) RNA-seq results of PHA-associated genes under 2% (w/v) supplemented NaCl stress relative to a control with 0.0% (w/v) supplemented NaCl, visualized as a dumbbell plot showing log₂ fold change (LFC) values. Genes are classified as downregulated (yellow), upregulated (purple), or not significantly differentially expressed (grey) (padj < 0.05 and |LFC| > 1).

Next, we analyzed the RNA-seq dataset to determine whether the observed changes in PHA content were associated with NaCl-responsive differential transcriptional expression of genes coding for the PHA biosynthetic machinery (**Figure 3B**; **Supplementary File S1**). Most PHA-related genes, including genes encoding acetyl-CoA acetyltransferase (*phaA*), acetoacetyl-CoA reductase (*phaB*), PHA synthases (*phaC1* and *phaC2*), PHA depolymerases (*phaZi* and *phaZe*) and PHA synthesis regulatory protein (*phaR*), were not significantly differentially expressed under osmotic stress (**Figure 3B**). Only the phasin-encoding gene *phaP* displayed a significant downregulation. Thus, alterations in PHA accumulation are unlikely to be strongly driven by transcriptional regulation.

To further contextualize salt-induced changes in PHA accumulation, proteins associated with purified PHA granules were analyzed by mass spectrometry (**Supplementary Figures S10** and **S11**; **Supplementary Tables S4** and **S5**). This revealed that granule-associated proteins were not limited to canonical PHA biosynthetic proteins. One of the most abundant proteins, both in SDS-PAGE band analysis and in shotgun LC-MS/MS, was a Hsp20/alpha-crystallin family protein (QPC31807.1; 16.8 kDa), accounting for approximately 20% of the total detected granule-associated protein abundance, supporting a strong association of this protein family with the PHA granule fraction (**Supplementary Table S4**). Interestingly, transcription of the gene encoding Hsp20/alpha-crystallin (IS481_01045) was upregulated under 2% (w/v) NaCl (LFC = +3.158). Among the classical PHA-associated proteins, PhaP was the most abundant canonical granule protein and was detected as the dominant protein both in SDS-PAGE band analysis and in shotgun LC-MS/MS. The latter analysis also led to the detection of PhaC1, PhaZi, PhaC2 and PhaR, while PhaA and PhaB were present at lower abundance. Additional non-canonical proteins were detected, including porins, OmpA-family proteins, TonB-dependent receptors, DUF883-family proteins, a universal stress protein and pilin-related proteins (**Supplementary Table S4**). These proteins may represent additional granule-associated proteins or, alternatively, might be co-purified membrane-associated proteins.

### 3.4. Salt-induced osmotic stress induces transcription of trehalose-associated metabolic pathways

Next, we analyzed the RNA-seq dataset to determine whether osmotic stress is associated with NaCl-responsive differential transcriptional expression of trehalose-associated genes, given its established role as a compatible solute and osmoprotectant (Elbein et al. 2003). Based on gene annotations, the three major microbial trehalose biosynthetic pathways are encoded in *C. thermodepolymerans* (**Figure 4A**). While the classical *de novo* trehalose synthesis pathway catalyzed by OtsA and OtsB did not show differential expression under increased supplemented NaCl concentrations, there was a strong upregulation of glucan-associated pathways (**Figure 4B**; **Supplementary File S1**). This includes the TreY–TreZ route (*treY*: LFC = +1.31; *treZ*: LFC = +1.57) and significant activation of glucans building and breaking enzymes (*glgX*: LFC = +2.11; *glgB*: LFC = +2.29), indicating mobilization of stored carbohydrates. The most pronounced induction was observed in the TreS–GlgE pathway (*treS/maK*: LFC = +2.43; *glgE*: LFC = +2.68), highlighting its central role in redirecting carbon flux towards trehalose production. Together, these transcriptional patterns suggest that, under salinity stress, *C. thermodepolymerans* preferentially shifts towards trehalose production via recycling and conversion of α-glucan-linked intermediates rather than relying on *de novo* synthesis.

**Figure 4.**
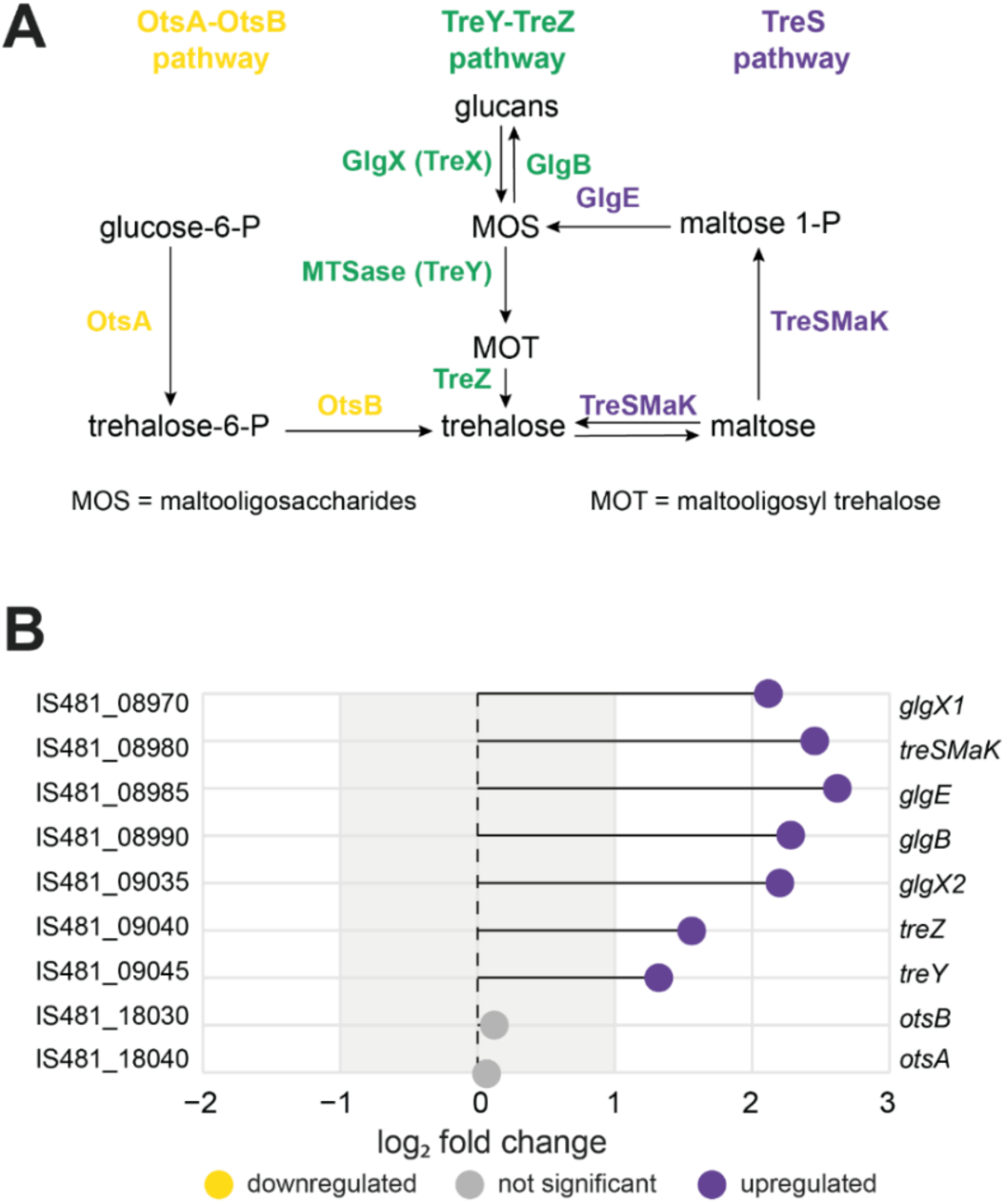
Trehalose biosynthetic pathways and transcriptional profiles of trehalose-associated genes in *C. thermodepolymerans* under NaCl stress. (**A**) Overview of the three major microbial trehalose biosynthetic pathways: the OtsA–OtsB pathway (yellow), the TreY–TreZ pathway (green) and the TreS pathway (purple). (**B**) RNA-seq results of trehalose-associated genes under 2% (w/v) supplemented NaCl stress relative to a control with 0.0% (w/v) supplemented NaCl, visualized as a dumbbell plot showing log₂ fold change values. Genes are classified as upregulated, downregulated or not significantly differentially expressed based on adjusted p-value < 0.05 and |LFC| > 1. The IS481_08980 gene is annotated as a fused trehalose synthase–maltokinase gene (*treSMaK*) (Urbániková and Janeček 2024).

### 3.5. Transcriptional upregulation of the type VI secretion system coincides with enhanced biofilm formation

The RNA-seq dataset revealed a prominent and coordinated transcriptional response of type VI secretion system (T6SS)-associated genes under 2% (w/v) supplemented NaCl (**Figure 5A**; **Supplementary File S1**). Several T6SS-related genes were strongly upregulated, with log₂ fold changes ranging from approximately 2 to 4.8, indicating a NaCl-responsive activation of this secretion-associated gene cluster under high-salt conditions.

**Figure 5.**
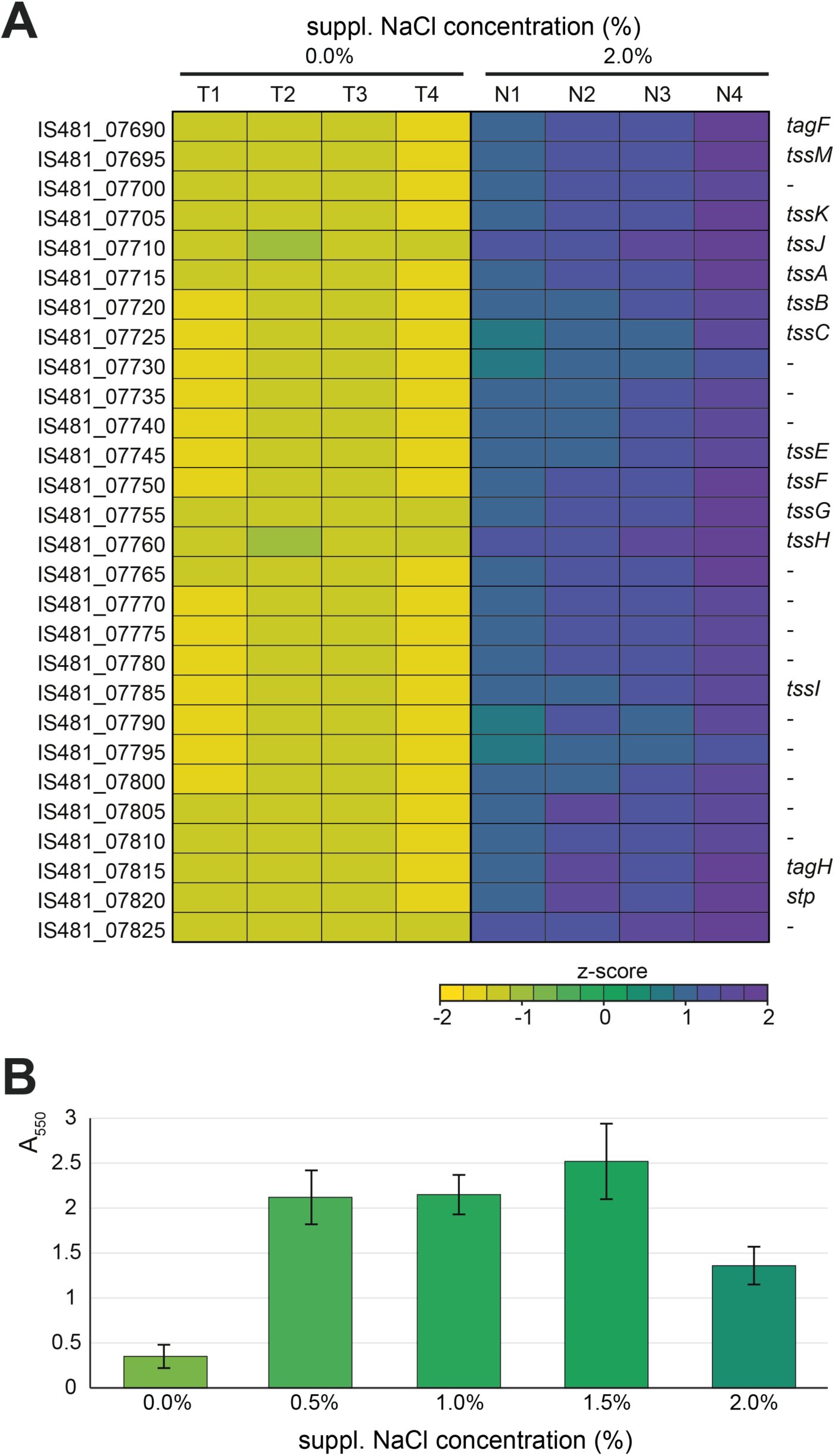
Transcriptional profiles of T6SS-associated genes and biofilm formation in *C. thermodepolymerans* under NaCl stress. (**A**) Transcriptional profiles of T6SS-associated genes under 2% (w/v) supplemented NaCl relative to a control in TSB medium without supplemented NaCl, visualized as a heatmap using z-scores calculated from normalized expression values for each gene. Columns represent biological replicates of the control condition (T1-T4; 0% supplemented NaCl) and NaCl stress condition (N1-N4; 2% supplemented NaCl). (**B**) Biofilm formation was assessed using a crystal violet assay after cultivation in TSB medium with different supplemented NaCl concentrations (0-2%, w/v). Absorbance was measured at 550 nm (A550), with five replicates per condition (n = 5) and Statistical significance was determined by one-way ANOVA with Tukey’s HSD test (*p* = 1.01E-08). Bars represent mean values and error bars indicate standard deviation.

Because T6SSs are commonly associated with intercellular interactions and surface-associated lifestyles, the transcriptional response of T6SS-associated genes prompted us to investigate biofilm formation. To this end, biofilm biomass was quantified using the crystal violet assay under increasing supplemented NaCl concentrations (**Figure 5B**; **Supplementary Figure S12**). Biofilm formation increased with NaCl supplementation, reaching the highest level at 1.5% (w/v) NaCl, where the absorption (A_550_) increased from 0.34 (control) to 2.45, corresponding to an approximately seven-fold increase. At 2% (w/v) NaCl, biofilm formation decreased again to an A₅₅₀ of 1.40, consistent with the reduced growth observed under this inhibitory NaCl concentration (**Figure 1**). Statistical analysis showed that biofilm formation increased significantly at 0.50%, 1.00%, and 1.50% NaCl compared with 0.00% NaCl, whereas 2.00% NaCl resulted in significantly lower biofilm formation than 0.50%, 1.00%, and 1.50% NaCl, while remaining significantly higher than the 0.00% NaCl control.

### 3.6. Salt-induced osmotic stress suppresses motility

The RNA-seq dataset indicated a global repression of genes associated with cell motility in salt-induced osmotic stress conditions, with the COG category N being enriched among downregulated genes (**Figure 2**). Indeed, the transcriptional profiles of most motility-associated genes displayed a significantly lower transcriptional expression in the presence of 2% (w/v) supplemented NaCl (**Figure 6A**; **Supplementary File S1**). This encompasses genes involved in flagellar assembly and function, including structural and regulatory flagellar genes (*flgA-K*, *fliA*, *fliC*, *fliE-T*, *flhA*, *flhB* and *flhD*), genes encoding the flagellar motor components (*motA* and *motB*), genes associated with pili- and surface-associated motility (*pilB*, *pilC*, *pilD*, *pilF*, *pilJ*, *pilL*, *pilM–P*, *pilT*, *pilW*, *pilY1* and *pilZ*) and type II secretion-associated genes (*gspD*, *gspE* and *gspE1*). Several chemotaxis-related genes (*cheA*, *cheB*, *cheD*, *cheW*, *cheY* and *cheZ*) also showed reduced expression.

**Figure 6.**
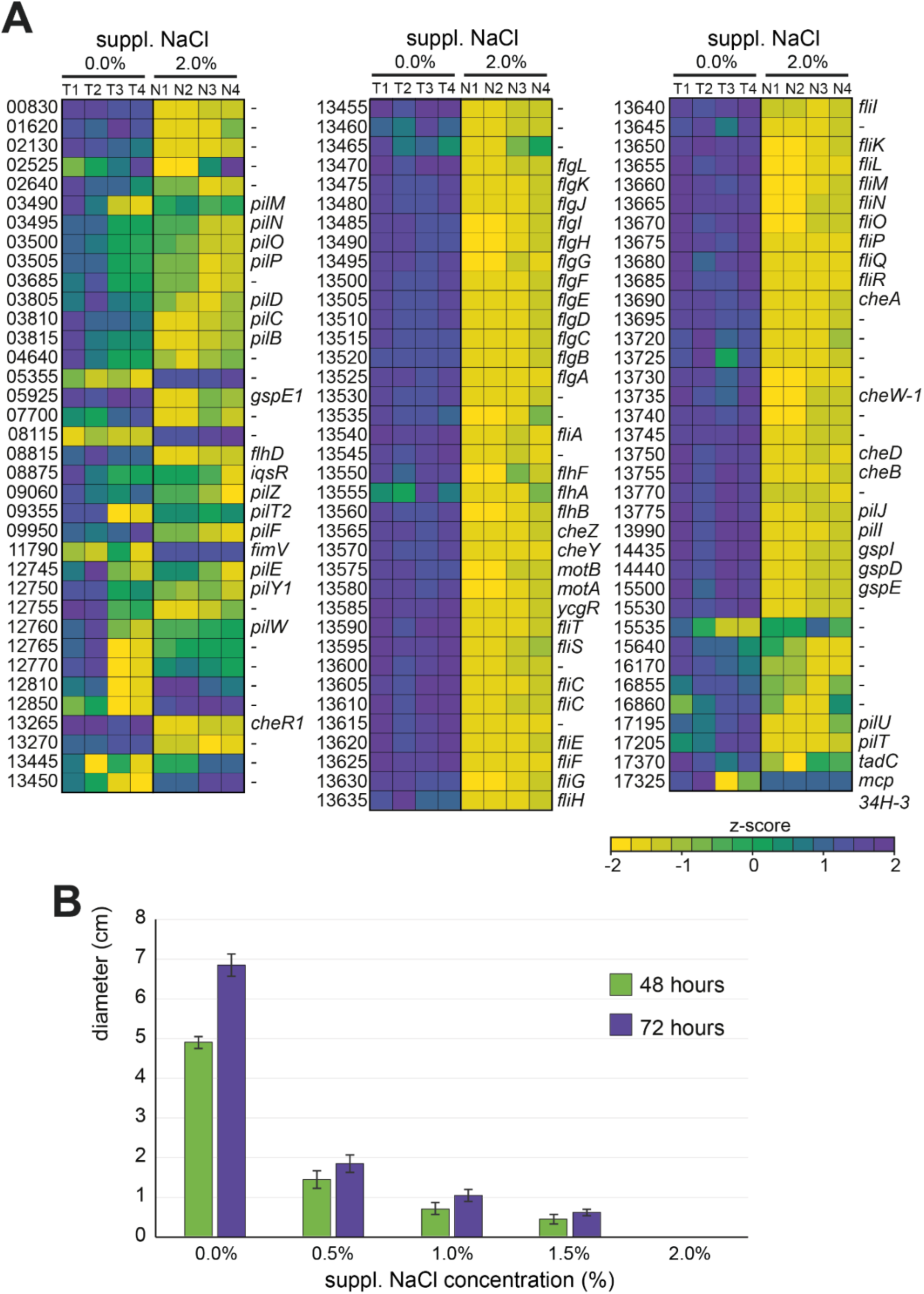
Transcriptional profiles of motility-associated genes and swarming motility of *C. thermodepolymerans* under NaCl stress. (**A**) Transcriptional profiles of flagellar, chemotaxis, pili and motility-associated genes under 2% (w/v) supplemented NaCl relative to a control in TSB medium without supplemented NaCl, visualized as a heatmap using z-scores calculated from normalized expression values for each gene. Columns represent biological replicates of the control condition (T1-T4; 0% supplemented NaCl) and the salt stress condition (N1-N4; 2% supplemented NaCl). (**B**) Swarming motility assessed on semi-solid TSA medium with supplemented NaCl (0-2% w/v) after 48 and 72 hours of incubation. Growth diameter was measured in biological triplicates (n = 3). Bars represent mean values and error bars indicate standard deviation.

The transcriptional repression of motility-associated genes was reflected phenotypically, as swarming motility was diminished when assessed on semi-solid medium with increasing supplemented NaCl concentrations (**Figure 6B**; **Supplementary Figure S13**). Swarming diameter decreased progressively with increasing NaCl concentration after both 48 and 72 hours of incubation. In unsupplemented medium, *C. thermodepolymerans* showed extensive surface spreading, whereas motility was strongly reduced at 0.5–1.5% (w/v) NaCl and was nearly abolished at 2% NaCl. Together, these results show that osmotic stress suppresses both motility-associated gene expression and the corresponding swarming phenotype in *C. thermodepolymerans*.

## 4. Discussion

This study demonstrates that salt-induced osmotic stress is a transcriptional and physiological driver in *C. thermodepolymerans*, linking salinity tolerance, PHA accumulation, osmoprotectant metabolism and surface-associated behavior. Moderate NaCl supplementation enhanced PHA accumulation, whereas 2% supplemented NaCl impaired growth and polymer accumulation, indicating that this condition approaches the upper physiological tolerance range of *C. thermodepolymerans*. This response is consistent with previous observations that salinity can affect PHA metabolism in a diverse range of prokaryotes, including non-halophilic PHA producers (Guzmán et al. 2009; Shrivastav et al. 2010; Cui et al. 2017; Godard et al. 2020; Obruca et al. 2021; Giovanella et al. 2025). These observations support the idea that salinity can act as a process-relevant lever for PHA production, but only within a defined stress window. Moreover, the increase in PHA accumulation under moderate salinity is compatible with the emerging view that PHA granules contribute not only to carbon and energy storage, but also to stress resistance (Obruca et al. 2018; 2020). In *C. necator*, PHA-containing cells showed reduced plasmolysis, membrane damage and cytoplasmic leakage under osmotic imbalance compared with PHA-deficient cells, suggesting that PHA granules can support cellular integrity during osmotic stress (Sedlacek et al. 2019).

The absence of broad transcriptional regulation of canonical *pha* genes, despite salt-associated changes in PHA accumulation, indicates that polymer accumulation under osmotic stress is unlikely to be controlled primarily at the level of transcription. Instead, it may involve regulation of carbon flux, enzyme activity, polymer turnover or granule organization (Hoffmann and Rehm 2004; Velázquez-Sánchez et al. 2020). This interpretation is supported by the PHA granule proteome, in which canonical granule-associated proteins such as PhaP and PhaC were detected, consistent with previous granule-proteome studies (Tirapelle et al. 2013), although Hsp20/alpha-crystallin family proteins were also highly abundant. A similar association of Hsp20 with PHA granules has been reported in *Pseudomonas extremaustralis* (Catone et al. 2014). This suggests a possible role for stress-responsive chaperone-like proteins in salinity-responsive PHA granule remodeling or stabilization (Laskowska et al. 2010; Dopson et al. 2017).

The transcriptional activation of trehalose-associated metabolism provides a second indication that salt-induced osmotic stress redirects carbon metabolism towards cellular protection. Trehalose is a well-established compatible solute that stabilizes proteins and membrane lipids under osmotic and dehydration stress (Elbein et al. 2003). In *C. thermodepolymerans*, the classical OtsA-OtsB pathway did not show differential expression, whereas glucan-linked routes were induced. These alternative routes have been reported to contribute to osmoadaptation and stress resilience in bacteria (Avonce et al. 2006; Chandra et al. 2011; Woodcock et al. 2021). This points to a broader metabolic strategy in which carbohydrate-derived osmoprotectants are mobilized to buffer salt-induced stress.

In parallel with intracellular osmoprotection, high NaCl concentrations appeared to promote a shift towards surface-associated behavior. The upregulation of T6SS-associated genes is notable because T6SSs are involved in bacterial interactions and have previously been linked to biofilm formation in several bacterial systems (Coulthurst 2019; Chen et al. 2020; Lin et al. 2021; Yang et al. 2022). In *C. thermodepolymerans*, upregulation of T6SS-associated genes coincided with enhanced biofilm formation at moderate NaCl concentrations, suggesting that osmotic stress favors surface-associated growth. Similar osmotic-stress-associated changes in biofilm formation have been reported in other bacteria, including *Clostridium ljungdahlii* and *Liberibacter crescens* (Philips et al. 2017; Padgett-Pagliai et al. 2022). Cell aggregation may also form part of the same surface-associated stress response, as aggregation has previously been described as an adaptation strategy under high-salinity conditions (Heinz et al. 2019). Although the presented results do not demonstrate a causal link between T6SS activity, aggregation and biofilm formation, they suggest that osmotic stress promotes community-level responses that may contribute to the protection of individual cells.

This surface-associated response was accompanied by a repression of motility-related functions, indicating a shift from active surface spreading towards a more sessile state. Such a shift is compatible with the concept that bacterial cells alter surface behaviors in response to environmental stress, although the direction and magnitude of motility responses are species-and condition-dependent. A similar stress-associated shift has been reported in *Clostridium ljungdahlii*, where high NaCl concentrations induced biofilm formation while flagellar motility was transcriptionally downregulated (Philips et al. 2017). Indeed, bacterial motility and biofilm formation are often reciprocally regulated, with repression of flagellar activity or flagellar gene transcription contributing to stabilization of surface-associated communities (Guttenplan and Kearns 2013; Dressaire et al. 2015). The repression of motility under NaCl-responsive stress is also consistent with reduced motility or flagellar activity reported in *Escherichia coli* BW25113, *Edwardsiella tarda*, *Clostridium ljungdahlii* and *Bacillus* sp. N16-5 under elevated salinity, where this response has been interpreted as part of ionic-stress adaptation or energy conservation (Steil et al. 2003; Yu et al. 2011; Yin et al. 2015; Philips et al. 2017; Li et al. 2021). Thus, reduced motility in *C. thermodepolymerans* may reflect both an energy-saving response and a transition towards surface-associated survival under osmotic stress.

Taken together, these observations support a model in which salt-induced osmotic stress redirects *C. thermodepolymerans* from growth-associated motility towards osmoprotection, carbon-storage-related processes and surface-associated persistence. From an applied perspective, this highlights moderate salt supplementation as a potential process lever for enhancing PHA accumulation, while also emphasizing that exceeding the physiological stress window compromises growth and polymer production.

## CRediT authorship contribution statement

**Mohamed Mostafa**: Conceptualization, Data curation, Formal analysis, Investigation, Software, Visualization, Writing – original draft. **Radwa Moanis**: Conceptualization, Data curation, Formal analysis, Investigation, Writing – review & editing. **Kristyna Hermankova**: Data curation, Formal analysis, Software, Visualization, Writing – review & editing. **Yannick Gansemans**: Data curation, Formal analysis, Software, Writing – review & editing. **Rani Baes**: Supervision, Writing – review & editing. **Filip Van Nieuwerburgh**: Resources, Writing – review & editing. **Karel Sedlar**: Supervision, Validation, Writing – review & editing. **Eveline Peeters**: Conceptualization, Funding acquisition, Supervision, Validation, Writing – review & editing.

## Acknowledgments

We are grateful to Karl Jonckheere for technical assistance and to Hannelore Geeraert and Niko Van den Brande for chemical characterization.

## Data availability

Sequencing data have been deposited in the NCBI Sequence Read Archive (SRA) under the project accession number PRJNA674774. SRA accession numbers for RNA-Seq samples are listed in **Supplementary Table S1** (SRR37584274 - SRR37584281). All other raw data underlying this study are available in either Supplementary Materials or a dataset with DOI 10.5281/zenodo.20571661. The code used for growth curve modelling is available at: https://github.com/MICR-VUB/RTS-8-Baranyi-Curve-fit-5-curves/.

## AI declaration

During the preparation of this work, OpenAI ChatGPT Plus was used to assist in polishing writing and debugging R code for fitting bacterial growth curves using the Baranyi growth model. After using this tool/service, the authors reviewed and edited the content as needed and take full responsibility for the content of the article.

## References

1. Adams, Jeremy David, Kyle B. Sander, Craig S. Criddle, Adam P. Arkin, and Douglas S. Clark. 2023. “Engineering osmolysis susceptibility in *Cupriavidus necator* and *Escherichia coli* for recovery of intracellular products.” Microbial Cell Factories 22 (1): 69. 10.1186/s12934-023-02064-8.

2. Andrews, S. 2010. “FastQC: a quality control tool for high throughput sequence data.” http://www.bioinformatics.babraham.ac.uk/projects/fastqc.

3. Avonce, Nelson, Alfredo Mendoza-Vargas, Enrique Morett, and Gabriel Iturriaga. 2006. “Insights on the evolution of trehalose biosynthesis.” BMC Evolutionary Biology 6 (1): 109. 10.1186/1471-2148-6-109.

4. Baranyi, József, and Terry A. Roberts. 1994. “A dynamic approach to predicting bacterial growth in food.” International Journal of Food Microbiology 23 (3–4): 277–94. 10.1016/0168-1605(94)90157-0.

5. Barik, Amita, and Santasabuj Das. 2018. “A comparative study of sequence- and structure-based features of small RNAs and other RNAs of bacteria.” RNA Biology 15 (1): 95–103. 10.1080/15476286.2017.1387709.

6. Behrends, V., J.G. Bundy, and H.D. Williams. 2011. “Differences in strategies to combat osmotic stress in *Burkholderia cenocepacia* elucidated by NMR-based metabolic profiling.” Letters in Applied Microbiology 52 (6): 619–25. 10.1111/j.1472-765X.2011.03050.x.

7. Bolger, Anthony M., Marc Lohse, and Bjoern Usadel. 2014. “Trimmomatic: a flexible trimmer for Illumina sequence data.” Bioinformatics 30 (15): 2114–20. 10.1093/bioinformatics/btu170.

8. Bosma, Elleke F., John Van Der Oost, Willem M. De Vos, and Richard Van Kranenburg. 2013. “Sustainable Production of Bio-Based Chemicals by Extremophiles.” Current Biotechnology 2 (4): 360–79. 10.2174/18722083113076660028.

9. Catone, Mariela V., Jimena A. Ruiz, Mildred Castellanos, Daniel Segura, Guadalupe Espin, and Nancy I. López. 2014. “High Polyhydroxybutyrate Production in *Pseudomonas extremaustralis* Is Associated with Differential Expression of Horizontally Acquired and Core Genome Polyhydroxyalkanoate Synthase Genes.” PLoS ONE 9 (6): e98873. 10.1371/journal.pone.0098873.

10. Chandra, Govind, Keith F. Chater, and Stephen Bornemann. 2011. “Unexpected and widespread connections between bacterial glycogen and trehalose metabolism.” Microbiology 157 (6): 1565–72. 10.1099/mic.0.044263-0.

11. Chen, Lihua, Yaru Zou, Asmaa Abbas Kronfl, and Yong Wu. 2020. “Type VI secretion system of *Pseudomonas aeruginosa* is associated with biofilm formation but not environmental adaptation.” MicrobiologyOpen 9 (3): e991. 10.1002/mbo3.991.

12. Chen, Shifu. 2025. “fastp 1.0: An ultra-fast all-round tool for FASTQ data quality control and preprocessing.” iMeta 4 (5): e70078. 10.1002/imt2.70078.

13. Coulthurst, Sarah. 2019. “The Type VI secretion system: a versatile bacterial weapon.” Microbiology 165 (5): 503–15. 10.1099/mic.0.000789.

14. Cox, Jürgen, and Matthias Mann. 2008. “MaxQuant enables high peptide identification rates, individualized p.p.b.-range mass accuracies and proteome-wide protein quantification.” Nature Biotechnology 26 (12): 1367–72. 10.1038/nbt.1511.

15. Cui, You-Wei, Xiao-Yu Gong, Yun-Peng Shi, and Zhiwu (Drew) Wang. 2017. “Salinity effect on production of PHA and EPS by *Haloferax mediterranei*.” RSC Advances 7 (84): 53587–95. 10.1039/C7RA09652F.

16. Dobin, Alexander, Carrie A. Davis, Felix Schlesinger, Jorg Drenkow, Chris Zaleski, Sonali Jha, Philippe Batut, Mark Chaisson, and Thomas R. Gingeras. 2013. “STAR: ultrafast universal RNA-seq aligner.” Bioinformatics 29 (1): 15–21. 10.1093/bioinformatics/bts635.

17. Dopson, Mark, David S. Holmes, Marcelo Lazcano, Timothy J. McCredden, Christopher G. Bryan, Kieran T. Mulroney, Robert Steuart, Connie Jackaman, and Elizabeth L. J. Watkin. 2017. “Multiple Osmotic Stress Responses in *Acidihalobacter prosperus* Result in Tolerance to Chloride Ions.” Frontiers in Microbiology 7 (January). 10.3389/fmicb.2016.02132.

18. Dressaire, Clémentine, Ricardo Neves Moreira, Susana Barahona, António Pedro Alves De Matos, and Cecília Maria Arraiano. 2015. “BolA Is a Transcriptional Switch That Turns Off Motility and Turns On Biofilm Development.” Edited by Susan Gottesman. mBio 6 (1): e02352–14. 10.1128/mBio.02352-14.

19. Elbanna, Khaled, Tina Lütke-Eversloh, Stefanie Van Trappen, Joris Mergaert, Jean Swings, and Alexander Steinbüchel. 2003. “*Schlegelella thermodepolymerans* gen. Nov., sp. Nov., a novel thermophilic bacterium that degrades poly(3-hydroxybutyrate-co-3-mercaptopropionate).” International Journal of Systematic and Evolutionary Microbiology 53 (4): 1165–68. 10.1099/ijs.0.02562-0.

20. Elbein, Alan D., Y.T. Pan, Irena Pastuszak, and David Carroll. 2003. “New insights on trehalose: a multifunctional molecule.” Glycobiology 13 (4): 17R–27R. 10.1093/glycob/cwg047.

21. Ewels, Philip, Måns Magnusson, Sverker Lundin, and Max Käller. 2016. “MultiQC: summarize analysis results for multiple tools and samples in a single report.” Bioinformatics 32 (19): 3047–48. 10.1093/bioinformatics/btw354.

22. Getino, Luis, José Luis Martín, and Alejandro Chamizo-Ampudia. 2024. “A Review of Polyhydroxyalkanoates: Characterization, Production, and Application from Waste.” Microorganisms 12 (10): 2028. 10.3390/microorganisms12102028.

23. Giovanella, Alice, Mónica Carvalheira, Matteo Grana, Maria A.M. Reis, and Bruno C. Marreiros. 2025. “Impact of saline osmotic stress on halotolerant polyhydroxyalkanoate (PHA)-accumulating mixed microbial cultures: Boosting PHA production by osmotic downshock.” Journal of Environmental Chemical Engineering 13 (3): 116521. 10.1016/j.jece.2025.116521.

24. Godard, Thibault, Daniela Zühlke, Georg Richter, Melanie Wall, Manfred Rohde, Katharina Riedel, Ignacio Poblete-Castro, Rainer Krull, and Rebekka Biedendieck. 2020. “Metabolic Rearrangements Causing Elevated Proline and Polyhydroxybutyrate Accumulation During the Osmotic Adaptation Response of *Bacillus megaterium*.” Frontiers in Bioengineering and Biotechnology 8 (February):47. 10.3389/fbioe.2020.00047.

25. Grybchuk-Ieremenko, Anastasiia, Kristýna Lipovská, Xenie Kouřilová, Stanislav Obruča, and Pavel Dvořák. 2025. “An Initial Genome Editing Toolset for *Caldimonas thermodepolymerans*, the First Model of Thermophilic Polyhydroxyalkanoates Producer.” Microbial Biotechnology 18 (2): e70103. 10.1111/1751-7915.70103.

26. Gu, Zuguang. 2022. “Complex heatmap visualization.” iMeta 1 (3): e43. 10.1002/imt2.43.

27. Guttenplan, Sarah B., and Daniel B. Kearns. 2013. “Regulation of flagellar motility during biofilm formation.” FEMS Microbiology Reviews 37 (6): 849–71. 10.1111/1574-6976.12018.

28. Guzmán, Héctor, Doan Van-Thuoc, Javier Martín, Rajni Hatti-Kaul, and Jorge Quillaguamán. 2009. “A process for the production of ectoine and poly(3-hydroxybutyrate) by *Halomonas boliviensis*.” Applied Microbiology and Biotechnology 84 (6): 1069–77. 10.1007/s00253-009-2036-2.

29. Heinz, Jacob, Annemiek C. Waajen, Alessandro Airo, Armando Alibrandi, Janosch Schirmack, and Dirk Schulze-Makuch. 2019. “Bacterial Growth in Chloride and Perchlorate Brines: Halotolerances and Salt Stress Responses of *Planococcus halocryophilus*.” Astrobiology 19 (11): 1377–87. 10.1089/ast.2019.2069.

30. Hoffmann, Nils, and Bernd H.A Rehm. 2004. “Regulation of polyhydroxyalkanoate biosynthesis in *Pseudomonas putida* and *Pseudomonas aeruginosa*.” FEMS Microbiology Letters 237 (1): 1–7. 10.1111/j.1574-6968.2004.tb09671.x.

31. Hrabalová, Vendula, Tomáš Opial, Jana Musilová, Karel Sedlář, and Stanislav Obruča. 2024. “Biotransformation of ferulic acid into vanillyl alcohol and vanillic acid employing thermophilic bacterium *Caldimonas thermodepolymerans*.” Enzyme and Microbial Technology 179 (September):110475. 10.1016/j.enzmictec.2024.110475.

32. Jang, Jun Won, In Yeub Hwang, Ok Kyung Lee, and Eun Yeol Lee. 2025. “Production of polyhydroxybutyrate with high cell density cultivation using thermophile *Caldimonas thermodepolymerans*.” Bioresource Technology 419 (March):132073. 10.1016/j.biortech.2025.132073.

33. Johnston, Brian, Iza Radecka, David Hill, Emo Chiellini, Vassilka Ivanova Ilieva, Wanda Sikorska, Marta Musioł, Magdalena Zięba, Adam A. Marek, Daniel Keddie, Barbara Mendrek, Surila Darbar, Grazyna Adamus, and Marek Kowalczuk. 2018. “The Microbial Production of Polyhydroxyalkanoates from Waste Polystyrene Fragments Attained Using Oxidative Degradation.” Polymers 10 (9): 957. 10.3390/polym10090957.

34. Koller, Martin. 2017. “Production of Polyhydroxyalkanoate (PHA) Biopolyesters by Extremophiles?” MOJ Polymer Science 1 (2). 10.15406/mojps.2017.01.00011.

35. Kopylova, Evguenia, Laurent Noé, and Hélène Touzet. 2012. “SortMeRNA: fast and accurate filtering of ribosomal RNAs in metatranscriptomic data.” Bioinformatics 28 (24): 3211–17. 10.1093/bioinformatics/bts611.

36. Kourilova, Xenie, Iva Pernicova, Karel Sedlar, Jana Musilova, Petr Sedlacek, Michal Kalina, Martin Koller, and Stanislav Obruca. 2020. “Production of polyhydroxyalkanoates (PHA) by a thermophilic strain of Schlegelella thermodepolymerans from xylose rich substrates.” Bioresource Technology 315 (November):123885. 10.1016/j.biortech.2020.123885.

37. Laskowska, Ewa, Ewelina Matuszewska, and Dorota Kuczynska-Wisnik. 2010. “Small Heat Shock Proteins and Protein-Misfolding Diseases.” Current Pharmaceutical Biotechnology 11 (2): 146–57. 10.2174/138920110790909669.

38. Li, Fen, Xue-Song Xiong, Ying-Ying Yang, Jun-Jiao Wang, Meng-Meng Wang, Jia-Wei Tang, Qing-Hua Liu, Liang Wang, and Bing Gu. 2021. “Effects of NaCl Concentrations on Growth Patterns, Phenotypes Associated With Virulence, and Energy Metabolism in *Escherichia coli* BW25113.” Frontiers in Microbiology 12 (August):705326. 10.3389/fmicb.2021.705326.

39. Li, Ru, Jian Yang, Yunzhu Xiao, and Lijuan Long. 2019. “In vivo immobilization of an organophosphorus hydrolyzing enzyme on bacterial polyhydroxyalkanoate nano-granules.” Microbial Cell Factories 18 (1): 166. 10.1186/s12934-019-1201-2.

40. Liao, Yang, Gordon K. Smyth, and Wei Shi. 2014. “featureCounts: an efficient general purpose program for assigning sequence reads to genomic features.” Bioinformatics 30 (7): 923–30. 10.1093/bioinformatics/btt656.

41. Lin, Jinshui, Lei Xu, Jianshe Yang, Zhuo Wang, and Xihui Shen. 2021. “Beyond dueling: roles of the type VI secretion system in microbiome modulation, pathogenesis and stress resistance.” Stress Biology 1 (1): 11. 10.1007/s44154-021-00008-z.

42. Love, Michael I, Wolfgang Huber, and Simon Anders. 2014. “Moderated estimation of fold change and dispersion for RNA-seq data with DESeq2.” Genome Biology 15 (12): 550. 10.1186/s13059-014-0550-8.

43. Mozejko-Ciesielska, Justyna, Dorota Dabrowska, Agnieszka Szalewska-Palasz, and Slawomir Ciesielski. 2017. “Medium-chain-length polyhydroxyalkanoates synthesis by *Pseudomonas putida* KT2440 relA/spoT mutant: bioprocess characterization and transcriptome analysis.” AMB Express 7 (1): 92. 10.1186/s13568-017-0396-z.

44. Musilova, Jana, Xenie Kourilova, Matej Bezdicek, Martina Lengerova, Stanislav Obruca, Helena Skutkova, and Karel Sedlar. 2021. “First Complete Genome of the Thermophilic Polyhydroxyalkanoates-Producing Bacterium *Schlegelella thermodepolymerans* DSM 15344.” Edited by Howard Ochman. Genome Biology and Evolution 13 (4): evab007. 10.1093/gbe/evab007.

45. Musilova, Jana, Xenie Kourilova, Kristyna Hermankova, Matej Bezdicek, Anastasiia Ieremenko, Pavel Dvorak, Stanislav Obruca, and Karel Sedlar. 2023. “Genomic and phenotypic comparison of polyhydroxyalkanoates producing strains of genus *Caldimonas*/*Schlegelella*.” Computational and Structural Biotechnology Journal 21:5372–81. 10.1016/j.csbj.2023.10.051.

46. Obruca, Stanislav, Petr Sedlacek, and Martin Koller. 2021. “The underexplored role of diverse stress factors in microbial biopolymer synthesis.” Bioresource Technology 326 (April):124767. 10.1016/j.biortech.2021.124767.

47. Obruca, Stanislav, Petr Sedlacek, Martin Koller, Dan Kucera, and Iva Pernicova. 2018. “Involvement of polyhydroxyalkanoates in stress resistance of microbial cells: Biotechnological consequences and applications.” Biotechnology Advances 36 (3): 856–70. 10.1016/j.biotechadv.2017.12.006.

48. Obruca, Stanislav, Petr Sedlacek, Eva Slaninova, Ines Fritz, Christina Daffert, Katharina Meixner, Zuzana Sedrlova, and Martin Koller. 2020. “Novel unexpected functions of PHA granules.” Applied Microbiology and Biotechnology 104 (11): 4795–4810. 10.1007/s00253-020-10568-1.

49. O’Toole, George A. 2011. “Microtiter Dish Biofilm Formation Assay.” Journal of Visualized Experiments : JoVE, no. 47 (January), 2437. 10.3791/2437.

50. Ozturk, Abdullah Bilal, Xenie Kourilova, Iva Buchtikova, and Stanislav Obruca. 2025. “Techno-economic assessment of polyhydroxyalkanoates production from lignocellulosic biomass employing halophilic and thermophilic microbial platform: Effect of fermentation conditions and downstream operations.” Waste Management 203 (July):114887. 10.1016/j.wasman.2025.114887.

51. Padgett-Pagliai, Kaylie A., Fernando A. Pagliai, Danilo R. Da Silva, Christopher L. Gardner, Graciela L. Lorca, and Claudio F. Gonzalez. 2022. “Osmotic stress induces long-term biofilm survival in *Liberibacter crescens*.” BMC Microbiology 22 (1): 52. 10.1186/s12866-022-02453-w.

52. Palma, Victoria, María Soledad Gutiérrez, Orlando Vargas, Raghuveer Parthasarathy, and Paola Navarrete. 2022. “Methods to Evaluate Bacterial Motility and Its Role in Bacterial–Host Interactions.” Microorganisms 10 (3): 563. 10.3390/microorganisms10030563.

53. Passanha, Pearl, Gopal Kedia, Richard M. Dinsdale, Alan J. Guwy, and Sandra R. Esteves. 2014. “The use of NaCl addition for the improvement of polyhydroxyalkanoate production by *Cupriavidus necator*.” Bioresource Technology 163 (July):287–94. 10.1016/j.biortech.2014.04.068.

54. Philips, Jo, Korneel Rabaey, Derek R. Lovley, and Madeline Vargas. 2017. “Biofilm Formation by *Clostridium ljungdahlii* Is Induced by Sodium Chloride Stress: Experimental Evaluation and Transcriptome Analysis.” Edited by Shihui Yang. PLOS ONE 12 (1): e0170406. 10.1371/journal.pone.0170406.

55. Schwanhäusser, Björn, Dorothea Busse, Na Li, Gunnar Dittmar, Johannes Schuchhardt, Jana Wolf, Wei Chen, and Matthias Selbach. 2011. “Global quantification of mammalian gene expression control.” Nature 473 (7347): 337–42. 10.1038/nature10098.

56. Sedlacek, Petr, Eva Slaninova, Martin Koller, Jana Nebesarova, Ivana Marova, Vladislav Krzyzanek, and Stanislav Obruca. 2019. “PHA granules help bacterial cells to preserve cell integrity when exposed to sudden osmotic imbalances.” New Biotechnology 49 (March):129–36. 10.1016/j.nbt.2018.10.005.

57. Shrivastav, Anupama, Sanjiv K. Mishra, and Sandhya Mishra. 2010. “Polyhydroxyalkanoate (PHA) synthesis by *Spirulina subsalsa* from Gujarat coast of India.” International Journal of Biological Macromolecules 46 (2): 255–60. 10.1016/j.ijbiomac.2010.01.001.

58. Smith, James L., and Brian J. Dell. 1990. “Capability of Selective Media to Detect Heat-Injured *Shigella flexneri*.” Journal of Food Protection 53 (2): 141–44. 10.4315/0362-028X-53.2.141.

59. Steil, Leif, Tamara Hoffmann, Ina Budde, Uwe Völker, and Erhard Bremer. 2003. “Genome-Wide Transcriptional Profiling Analysis of Adaptation of *Bacillus subtilis* to High Salinity.” Journal of Bacteriology 185 (21): 6358–70. 10.1128/JB.185.21.6358-6370.2003.

60. Tirapelle, Evandro F., Marcelo Müller-Santos, Michelle Z. Tadra-Sfeir, Marco A. S. Kadowaki, Maria B. R. Steffens, Rose A. Monteiro, Emanuel M. Souza, Fabio O. Pedrosa, and Leda S. Chubatsu. 2013. “Identification of Proteins Associated with Polyhydroxybutyrate Granules from *Herbaspirillum seropedicae* SmR1 - Old Partners, New Players.” Edited by Mickaël Desvaux. PLoS ONE 8 (9): e75066. 10.1371/journal.pone.0075066.

61. Urbániková, Ľubica, and Štefan Janeček. 2024. “Trehalose synthases from the subfamily GH13_16 involved in α-glucan biosynthesis – a focus on their maltokinase domain.” International Journal of Biological Macromolecules 268 (May):131680. 10.1016/j.ijbiomac.2024.131680.

62. Velázquez-Sánchez, Claudia, Guadalupe Espín, Carlos Peña, and Daniel Segura. 2020. “The Modification of Regulatory Circuits Involved in the Control of Polyhydroxyalkanoates Metabolism to Improve Their Production.” Frontiers in Bioengineering and Biotechnology 8 (April):386. 10.3389/fbioe.2020.00386.

63. Wickham, Hadley. 2016. *ggplot2: Elegant Graphics for Data Analysis*. Use R! Cham: Springer International Publishing. 10.1007/978-3-319-24277-4.

64. Woodcock, Stuart D., Karl Syson, Richard H. Little, Danny Ward, Despoina Sifouna, James K. M. Brown, Stephen Bornemann, and Jacob G. Malone. 2021. “Trehalose and α-glucan mediate distinct abiotic stress responses in *Pseudomonas aeruginosa*.” PLOS Genetics 17 (4): e1009524. 10.1371/journal.pgen.1009524.

65. Yang, Yantao, Damin Pan, Yanan Tang, Jiali Li, Kaixiang Zhu, Zonglan Yu, Lingfang Zhu, Yao Wang, Peng Chen, and Changfu Li. 2022. “H3-T6SS of *Pseudomonas aeruginosa* PA14 contributes to environmental adaptation via secretion of a biofilm-promoting effector.” Stress Biology 2 (1): 55. 10.1007/s44154-022-00078-7.

66. Yin, Liang, Yanfen Xue, and Yanhe Ma. 2015. “Global Microarray Analysis of Alkaliphilic Halotolerant Bacterium *Bacillus* sp. N16-5 Salt Stress Adaptation.” PLOS ONE 10 (6): e0128649. 10.1371/journal.pone.0128649.

67. Yu, Jong-Earn, Park, Jun-Mo, and Kang, Ho Young. 2011. “Effects of Salt Concentration on Motility and Expression of Flagellin Genes in the Fish Pathogen *Edwardsiella tarda*.” Journal of Life Science 21 (10): 1487–93. 10.5352/JLS.2011.21.10.1487.

68. Zhou, Wen, Dana Irene Colpa, Hjalmar Permentier, Ruben Ate Offringa, Leon Rohrbach, Gert-Jan Willem Euverink, and Janneke Krooneman. 2023. “Insight into polyhydroxyalkanoate (PHA) production from xylose and extracellular PHA degradation by a thermophilic *Schlegelella thermodepolymerans*.” Resources, Conservation and Recycling 194 (July):107006. 10.1016/j.resconrec.2023.107006.

69. Zhou, Wen, Salvador Bertrán Llorens, Peter J. Deuss, Gert-Jan W. Euverink, and Janneke Krooneman. 2025. “Polyhydroxyalkanoate (PHA) production by thermophilic *Caldimonas thermodepolymerans* comb. nov. from xylan.” RSC Sustainability 3 (4): 1685–90. 10.1039/D5SU00040H.

