## Supplementary Materials for "Salt-induced osmotic stress remodels osmoadaptive gene expression and physiology in the polyhydroxyalkanoate-accumulating thermophilic bacterium *Caldimonas thermodepolymerans*"

Mostafa *et al*.

### Supplementary Figures

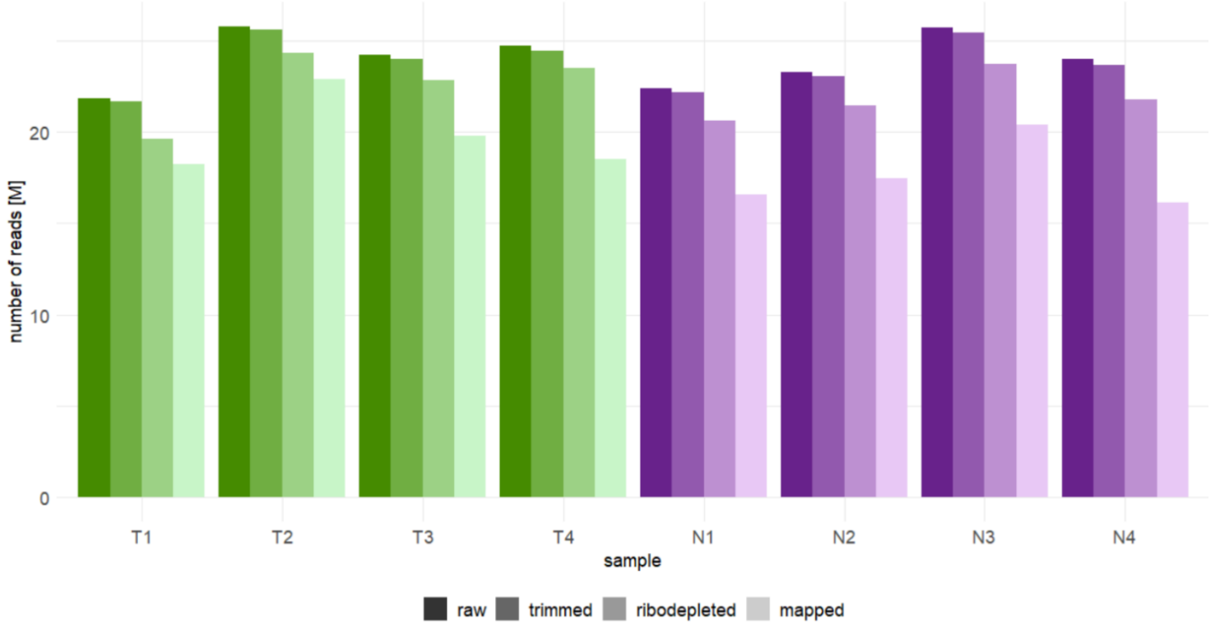


**Supplementary Figure S1.** Read counts of RNA-seq samples during data preprocessing.

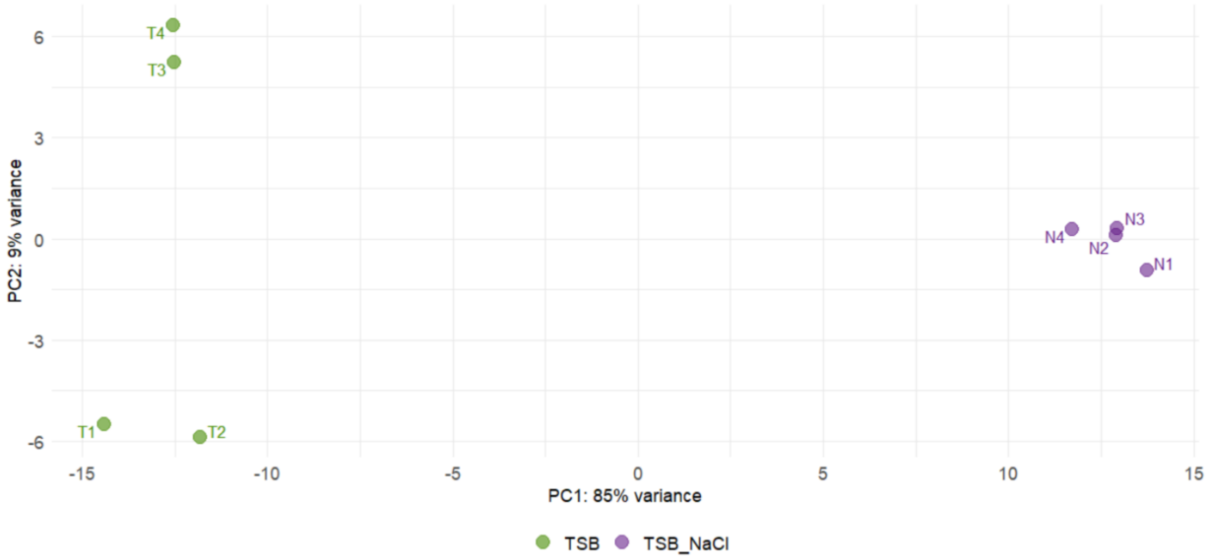


**Supplementary Figure S2.** Principal component analysis of RNA-seq samples.

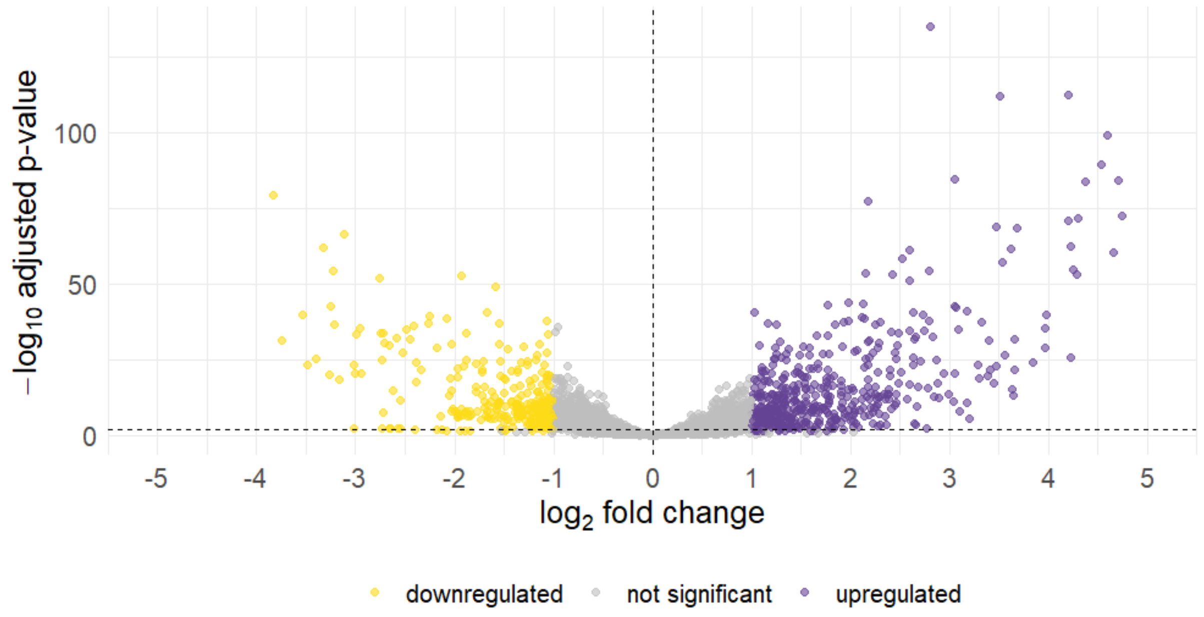


**Supplementary Figure S3**. Volcano plot of differentially expressed genes in *C. thermodepolymerans* grown under 2% supplemented NaCl as compared to a control without supplemented NaCl. Genes meeting adjusted p-value < 0.05 and $|\log_2$ fold change$| > 1$ are shown as upregulated (purple; n = 485) or downregulated (yellow; n = 279), while non-significant genes are shown in grey.

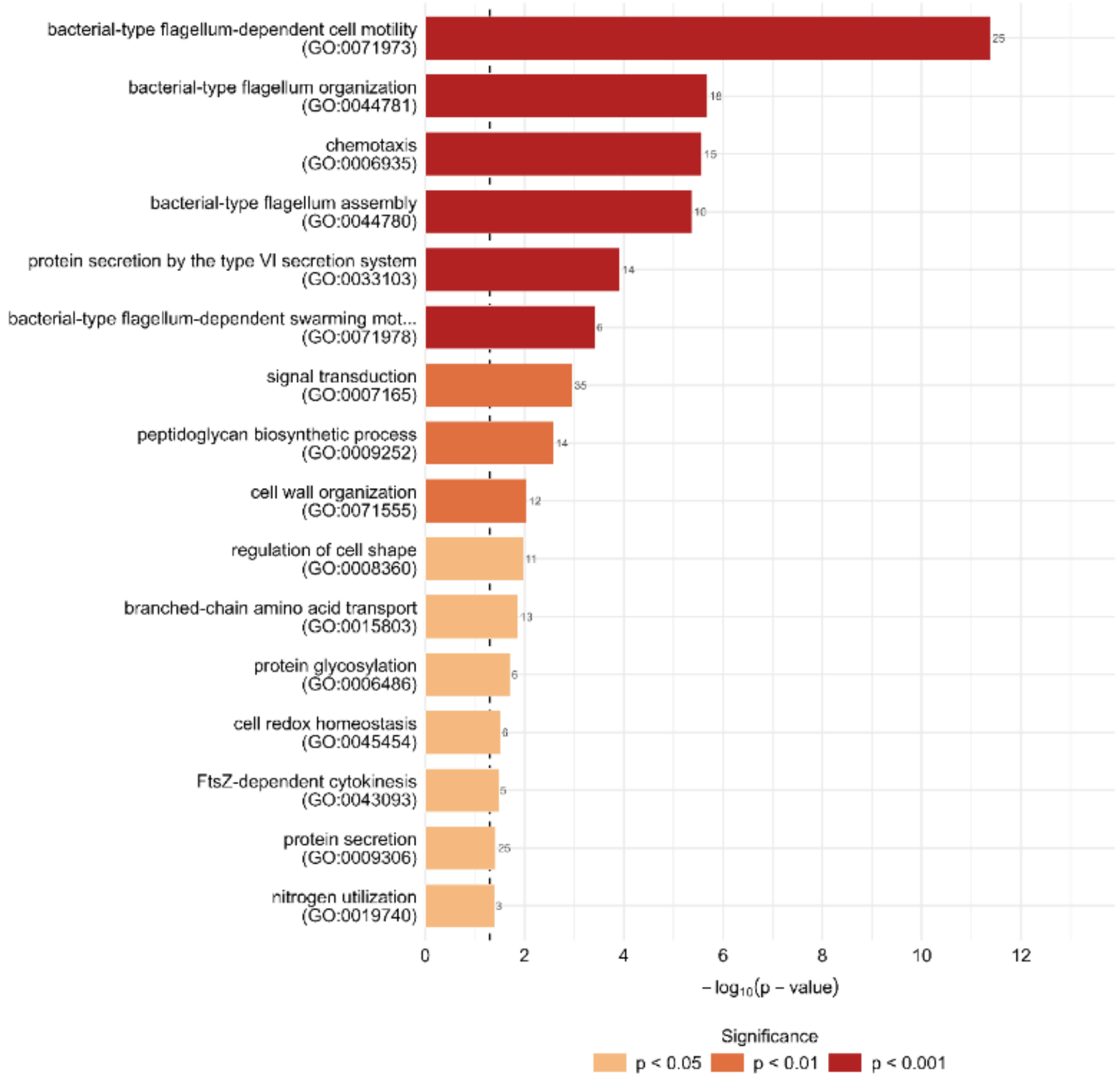


**Supplementary Figure S4.** Significantly enriched Gene Ontology (Biological Process) terms.

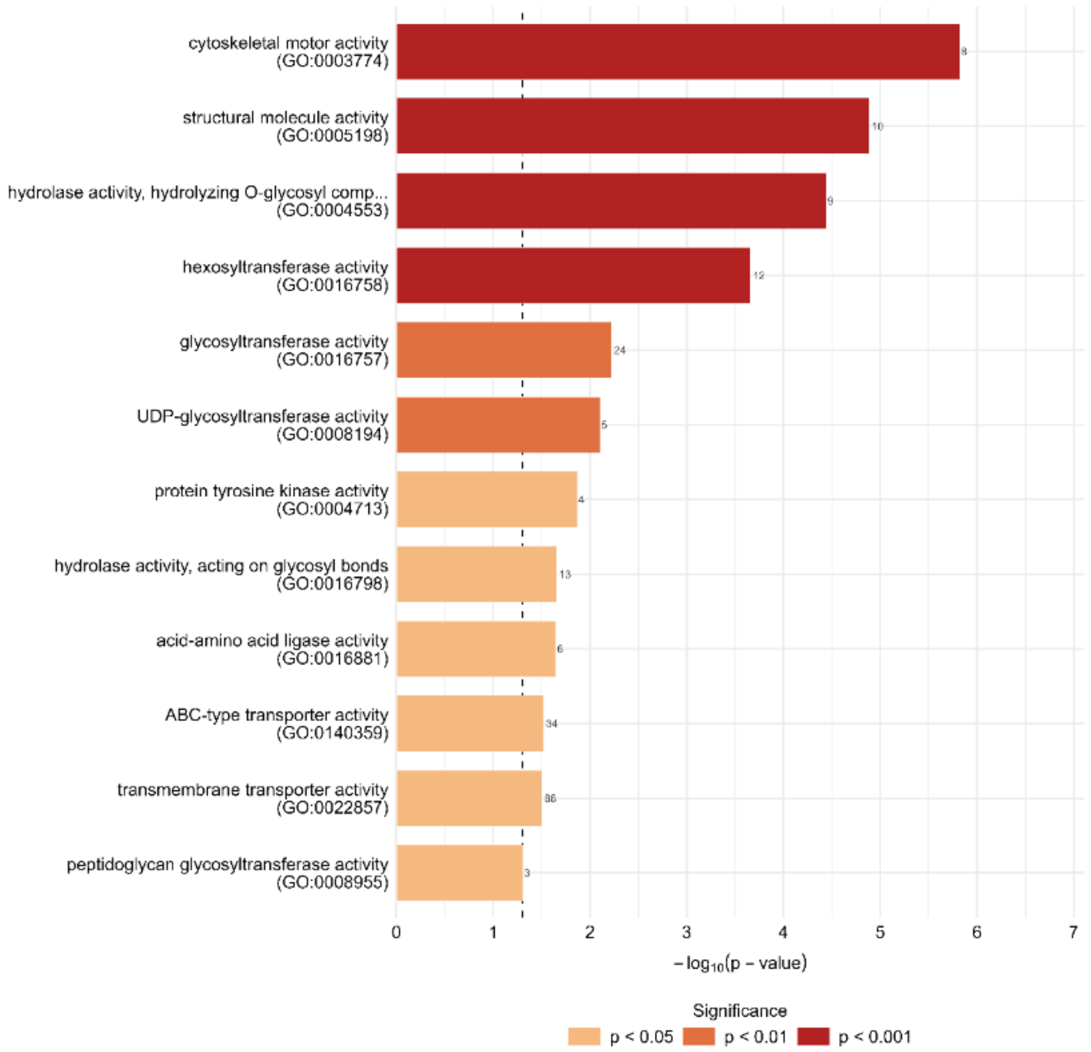


**Supplementary Figure S5.** Significantly enriched Gene Ontology (Molecular Function) terms.

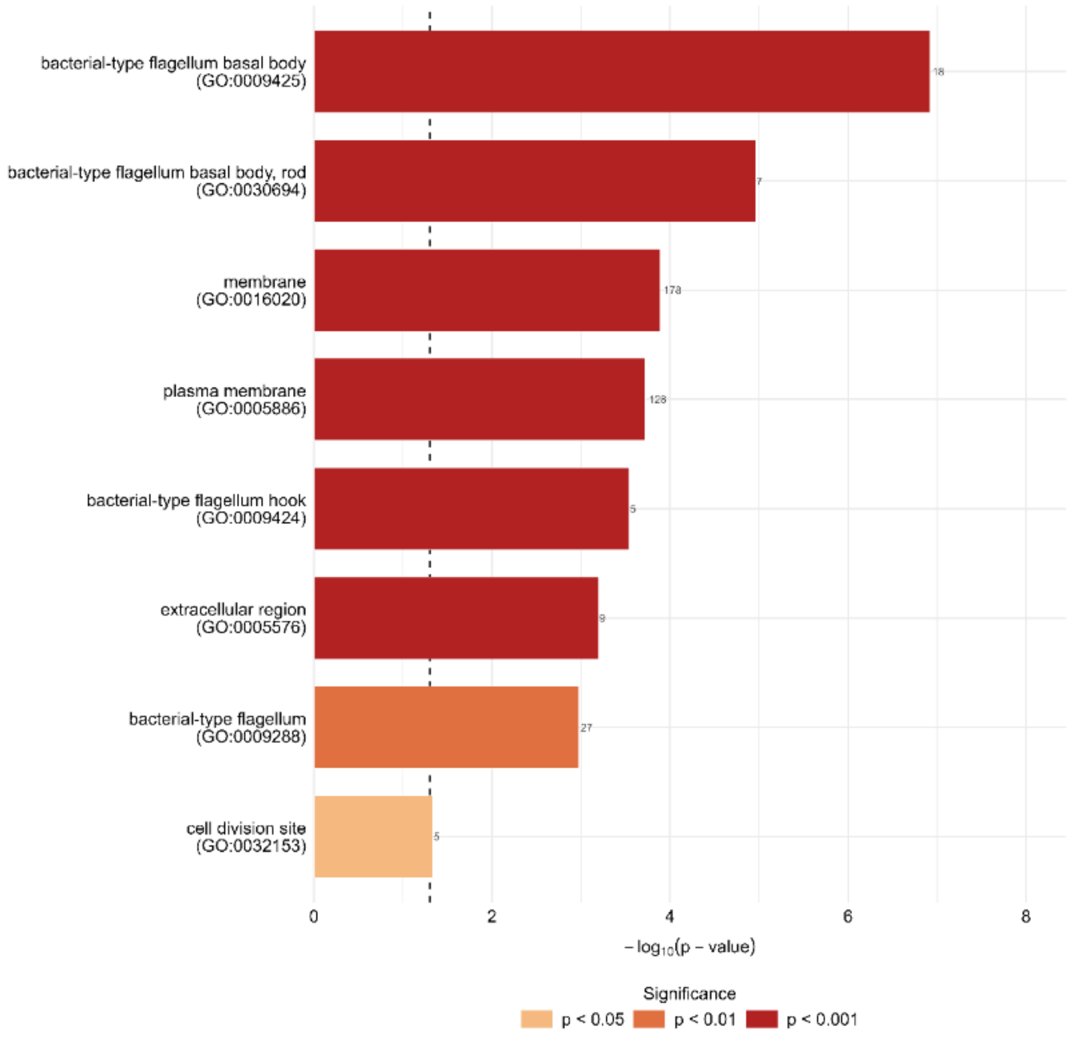


**Supplementary Figure S6.** Significantly enriched Gene Ontology (Cellular Component) terms.

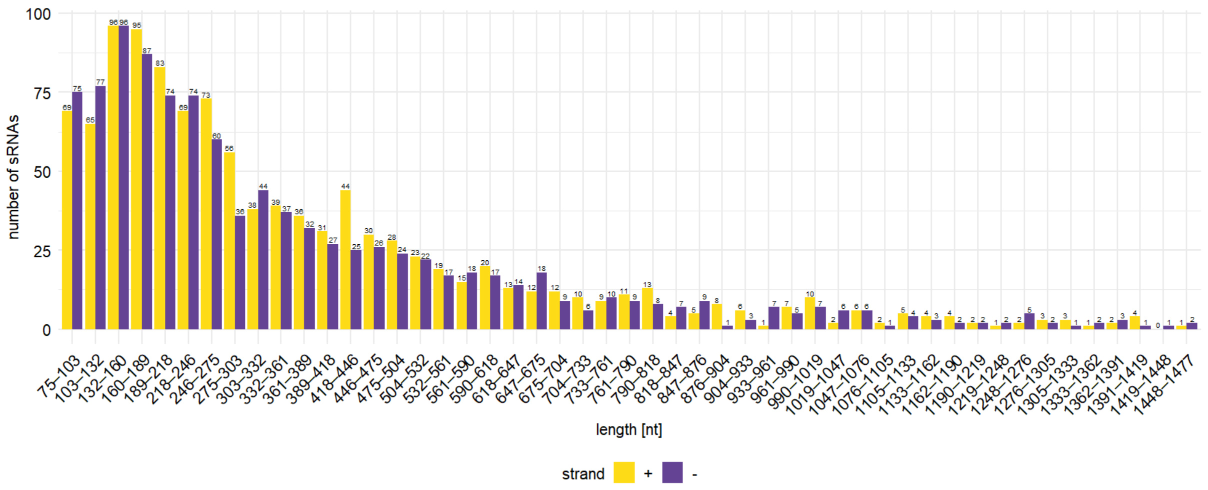


**Supplementary Figure S7.** Length distribution of putative small RNAs predicted in the genome of *C. thermodepolymerans* 15344$^{T}$ after length filtering (>1500 nt).

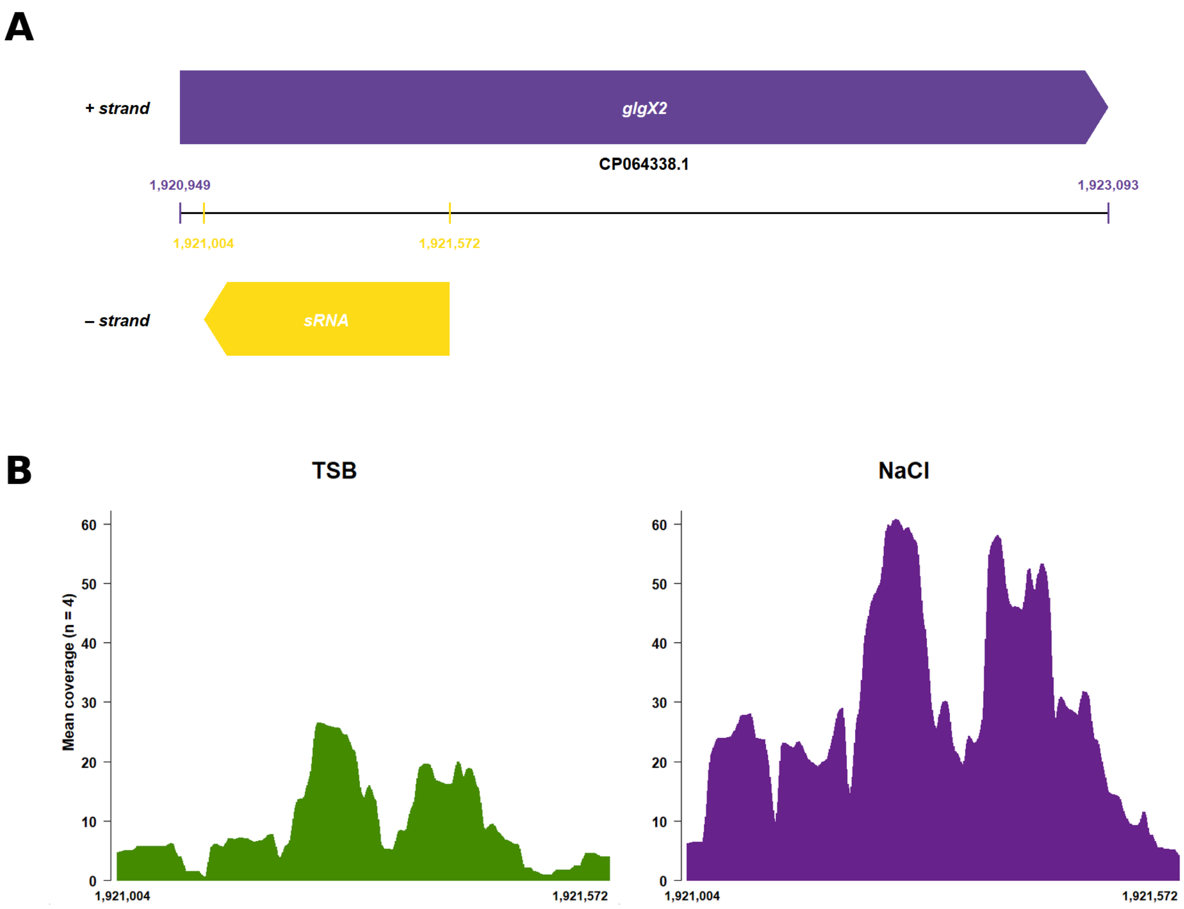


**Supplementary Figure S8. Example of a *cis*-encoded antisense sRNA. (A)** Schematic representation of the genomic location of the *cis*-encoded sRNA relative to *glgX2* (IS481_09035). The sRNA is transcribed from the opposite strand and overlaps the 5' region of the coding sequence. **(B)** Strand-specific RNA-seq coverage (mean of four biological replicates) showing increased expression of the *cis*-encoded sRNA under 2% NaCl compared with TSB. The overlapping *glgX2* transcript displays a similar expression trend (see Fig. 4B), suggesting coordinated expression under osmotic stress.

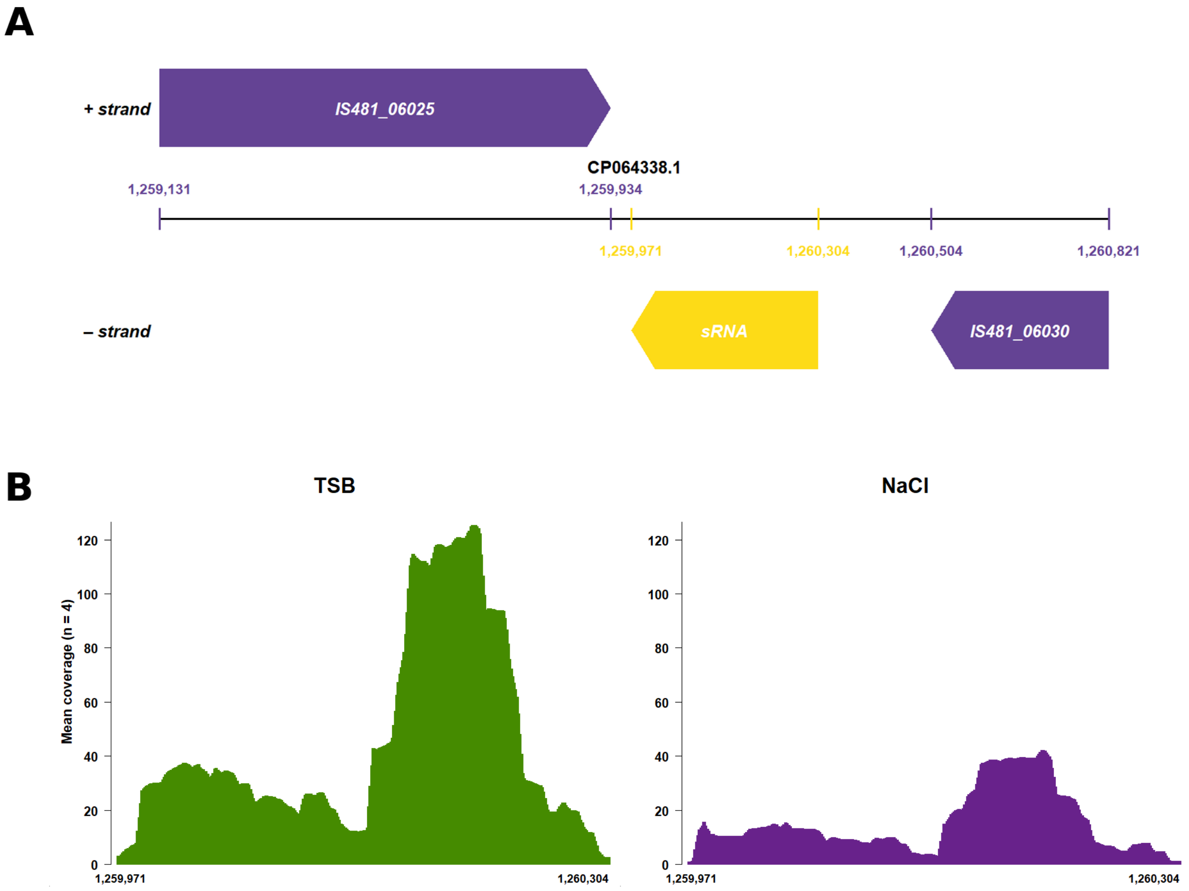


**Supplementary Figure S9. Example of a *trans*-encoded sRNA. (A)** Genomic location of the *trans*-encoded sRNA in the intergenic region between **IS481_06025** and **IS481_06030**. **(B)** Strand-specific RNA-seq coverage showing higher expression of the *trans*-encoded sRNA in TSB than under 2% NaCl conditions.

(A)

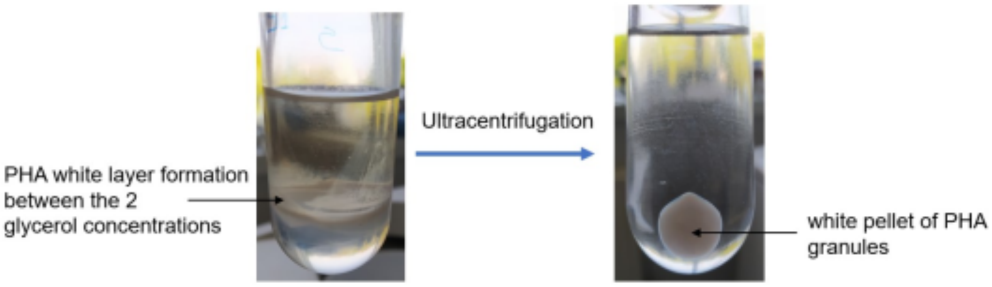


(B)

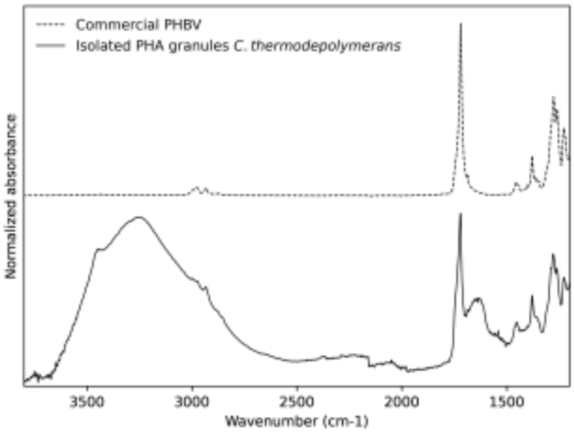


**Supplementary Figure S10.** (**A**) PHA granules extraction by glycerol-gradient ultracentrifugation showing the formation of PHA granules white layer between the 2 glycerol layers, which was then further ultracentrifuged to form a white pellet of PHA, (**B**) FTIR spectrum of the extracted PHA granules, commercial PHBV was analyzed as a reference.

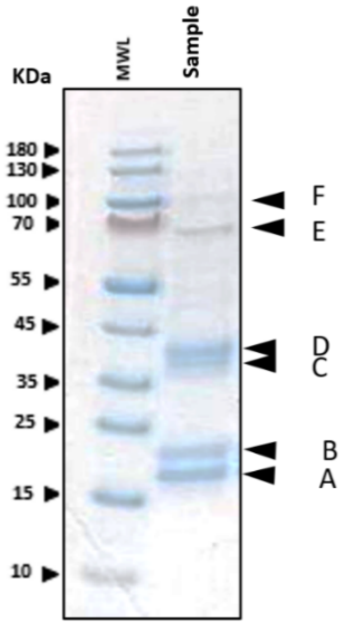


**Supplementary Figure S11.** SDS-PAGE showing the bands corresponding to the most abundant proteins attached to PHA granules, further analyzed by mass spectrometry (**Supplementary Table S4**). MWL: molecular weight ladder.

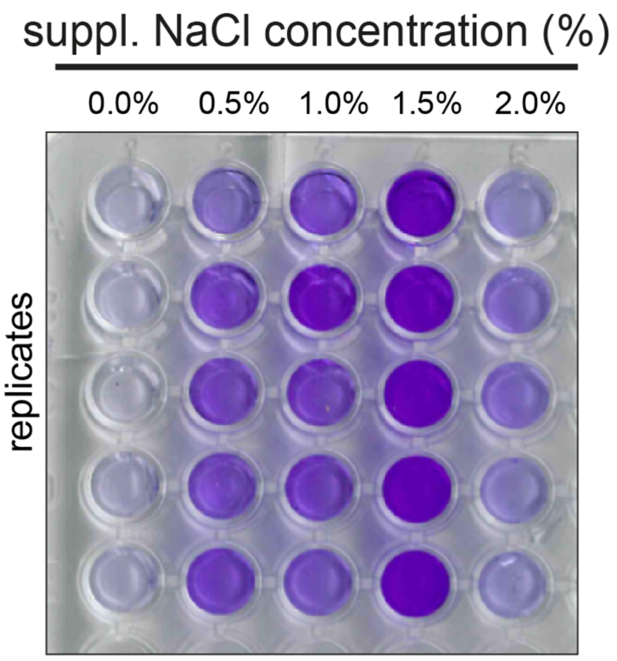


**Supplementary Figure S12**. Crystal-violet-stained microtiter plate used for biofilm quantification in *C. thermodepolymerans* under different supplemented NaCl (w/v) concentrations.

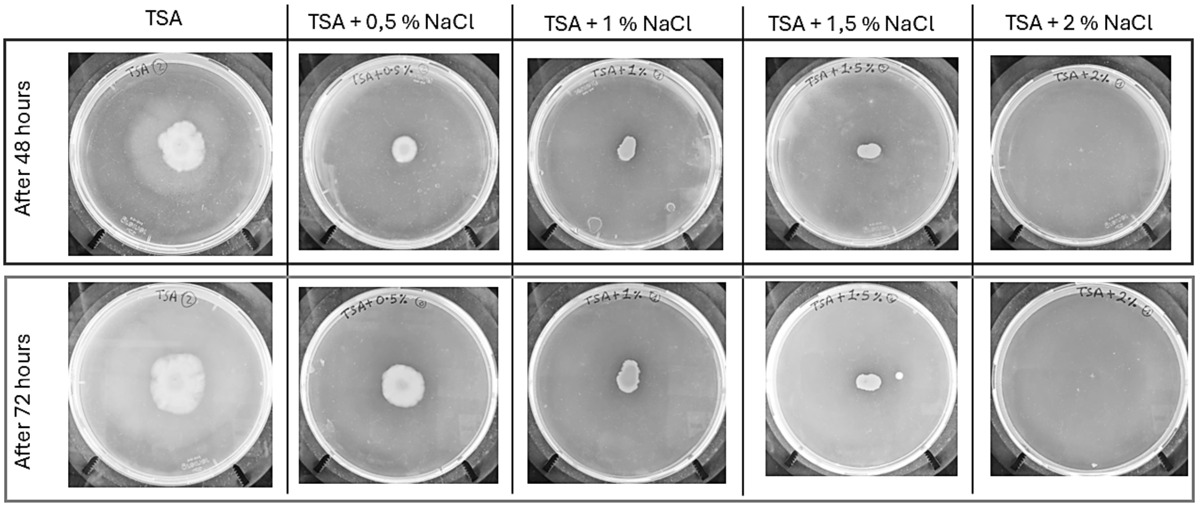


**Supplementary Figure S13.** Representative swarming motility plates of *C. thermodepolymerans* cultivated on TSA medium supplemented with increasing NaCl concentrations. Swarming motility was assessed by measuring the growth diameter across the widest spreading zone in biological triplicates (n = 3).

**Supplementary Tables**

**Supplementary Table S1**. Growth parameters of *C. thermodepolymerans* DSM 15344$^T$ cultivated in TSB medium supplemented with 0–2% NaCl. Values are ± Standard deviation (SD), and RMSE = root mean square error.

| **Condition** | **$y_0$ (initial $OD_{600}$)** | **$y_{max}$ (max $OD_{600}$)** | **μ ($h^{-1}$)** | **Lag time (h)** | **RMSE** |
|---|---|---|---|---|---|
| TSB(Control) | 0.472±0.005 | 1.301605±0.026 | 2.761±0.119 | 0.884±0.055 | 0.0325±0.0003 |
| TSB+0.5% NaCl | 0.538±0.009 | 1.35512±0.020 | 2.192±0.245 | 0.966±0.004 | 0.0386±0.004 |
| TSB+1.0% NaCl | 0.445±0.002 | 1.353245±0.001 | 2.074±0.004 | 1.447±0.001 | 0.0278±0.0001 |
| TSB+1.5% NaCl | 0.464±0.0007 | 1.23614±0.013 | 1.102±0.029 | 1.948±0.029 | 0.0411±0.0008 |
| TSB+2.0% NaCl | 0.476±0.002 | 0.916934±0.002 | 1.358±0.036 | 1.793±0.002 | 0.0303±0.0015 |

**Supplementary Table S2.** Overview of RNA-seq samples, quality control (QC) and mapped reads.

| **Sample name** | **Substrate** | **Replicate** | **SRA acc. no** | **Total number of reads [M]** | **Number of QC reads [M]** | **Number of non-rRNA reads [M]** | **Number of mapped reads [M]** |
|---|---|---|---|---|---|---|---|
| T1 | TSB | 1 | SRR37584281 | 21.83 | 21.66 | 19.63 | 18.24 |
| T2 | TSB | 2 | SRR37584280 | 25.8 | 25.59 | 24.34 | 22.89 |
| T3 | TSB | 3 | SRR37584279 | 24.24 | 24.02 | 22.81 | 19.8 |
| T4 | TSB | 4 | SRR37584278 | 24.74 | 24.45 | 23.48 | 18.51 |
| N1 | TSB+NaCl | 1 | SRR37584277 | 22.42 | 22.15 | 20.63 | 16.56 |
| N2 | TSB+NaCl | 2 | SRR37584276 | 23.29 | 23.04 | 21.45 | 17.46 |
| N3 | TSB+NaCl | 3 | SRR37584275 | 25.7 | 25.42 | 23.74 | 20.41 |

**Supplementary Table S3.** Clusters of orthologous groups (COG) class representation in the genome of *C. thermodepolymerans* DSM 15344$^T$.

| COG class | Class description | Gene count | % from all CDS |
|---|---|---|---|
| A | RNA processing and modification | 1 | 0.03 |
| B | Chromatin structure and dynamics | 2 | 0.06 |
| C | Energy production and conversion | 275 | 7.66 |
| D | Cell cycle control, cell division, chromosome partitioning | 39 | 1.09 |
| E | Amino acid transport and metabolism | 280 | 7.80 |
| F | Nucleotide transport and metabolism | 94 | 2.62 |
| G | Carbohydrate transport and metabolism | 126 | 3.51 |
| H | Coenzyme transport and metabolism | 148 | 4.12 |
| I | Lipid transport and metabolism | 160 | 4.46 |
| J | Translation, ribosomal structure and biogenesis | 178 | 4.96 |
| K | Transcription | 258 | 7.19 |
| L | Replication, recombination and repair | 150 | 4.18 |
| M | Cell wall/membrane/envelope biogenesis | 192 | 5.35 |
| N | Cell motility | 98 | 2.73 |
| O | Posttranslational modification, protein turnover, chaperones | 123 | 3.43 |
| P | Inorganic ion transport and metabolism | 168 | 4.68 |
| Q | Secondary metabolites biosynthesis, transport, and catabolism | 78 | 2.17 |
| S | Function unknown | 681 | 18.97 |
| T | Signal transduction mechanisms | 130 | 3.62 |
| U | Intracellular trafficking, secretion, and vesicular transport | 70 | 1.95 |
| V | Defense mechanisms | 29 | 0.81 |
| - | Unknown | 309 | 8.61 |
| **sum** | | **3589** | **100** |

**Supplementary Table S4.** The most abundant protein detected in the bands obtained from running Granule-attached proteins on SDS-PAGE. E*: the two most abundant proteins in the band are mentioned here. MW = molecular weight.

| **Band** | **Protein IDs** | **Protein name** | **Relative abundance in the band** | **MW (kDa)** |
|---|---|---|---|---|
| A | QPC31807.1 | Hsp20/alpha-crystallin family protein | 48% | 16.78 |
| B | QPC32978.1 | Phasin family protein (**PhaP2**) | 77% | 19.21 |
| C | QPC31677.1 | Porin | 48% | 33.02 |
| D | QPC31109.1 | Porin | 59% | 34.70 |
| E* | QPC30057.1 | TonB-dependent receptor | 69% | 61.14 |
| | QPC31672.1 | Polyhydroxyalkanoic acid synthase (**PhaC1**) | 6% | 65.09 |
| F | QPC31774.1 | TonB-dependent receptor | 36% | 82.28 |

**Supplementary Table S5.** All common PHA enzymes that were detected attached to PHA granules via proteomics analysis. Protein abundance is indicated by the iBAQ value, which corresponds to the sum of all peptide intensities divided by the number of observable peptides of a protein. MW = molecular weight.

| Identified PHA enzyme | Locus tags | Protein IDs | iBAQ value ($x10^3$) | MW (kDa) |
|---|---|---|---|---|
| Phasin (PhaP2) | IS481_07490 | QPC32978.1 | 19841000 | 19.24 |
| Polyhydroxyalkanoate synthase (PhaC1) | IS481_00330 | QPC31672.1 | 1032100 | 66.00 |
| Polyhydroxyalkanoate depolymerase (PhaZ) | IS481_07130 | QPC32911.1 | 762970 | 47.06 |
| Polyhydroxyalkanoate synthase (PhaC2) | IS481_08630 | QPC33188.1 | 417060 | 62.73 |
| polyhydroxyalkanoate synthesis repressor (PhaR) | IS481_08360 | QPC33138.1 | 63058 | 22.04 |
| acetyl-CoA C-acetyltransferase (PhaA) | IS481_08635 | QPC33189.1 | 20644 | 40.76 |
| acetoacetyl-CoA reductase (PhaB) | IS481_08640 | QPC33190.1 | 18361 | 26.27 |

### Supplementary Notes

#### Supplementary Note S1. RNA-sequencing data analysis.

##### *COG and GO enrichment*

For additional functional annotation, Clusters of Orthologous Groups (COG) assignments were obtained using EggNOG-mapper (v2.1.13) (Cantalapiedra et al. 2021). These annotations were subsequently used for COG enrichment analysis implemented in a custom R script. For this analysis, all genes tested for differential expression (DE) were used as the background gene set. Enrichment was evaluated separately for genes with positive (LFC > 0) and negative (LFC < 0) $\log_2$ fold changes within each COG category. Statistical significance was assessed using Fisher's exact test, and the resulting p-values were corrected for multiple testing, again using the BH method. Categories with padj < 0.05 were considered significantly enriched.

Gene Ontology (GO) terms were obtained from the QuickGO (Binns et al. 2009) annotation file for taxon 215580 (*Caldimonas thermodepolymerans*), together with a file containing all proteins associated with *C. thermodepolymerans* in the UniProt database (Ahmad et al. 2025). Using a custom R script, GO terms were mapped to locus tags via UniProt IDs and protein IDs of coding sequences (CDSs) extracted from the GenBank file. These annotations were then used to perform GO enrichment analysis using the topGO package (v2.54.0) (Alexa and Rahnenführer 2025). All genes tested for DE were used as the gene universum. Enrichment was conducted using the 'weight01' algorithm, which accounts for the hierarchical structure of GO and down-weights genes annotated to multiple terms, reducing bias from highly connected nodes. Statistical significance was assessed using Fisher's exact test. No multiple testing correction was applied in this analysis, as the 'weight01' algorithm already mitigates the impact of highly overlapping GO terms.

##### *Small RNA analysis*

The analysis of small RNA (sRNA) content began with sRNA prediction using the baerhunter package (v0.9.1.0) (Ozuna et al. 2020). Predicted sRNAs were subsequently filtered based on their length (<1500 nt). Further, the sRNAs were divided into *cis*/*trans*-encoded according to genomic overlap using the IRanges package (v2.36.0) (Lawrence et al. 2013). *Cis*-encoded sRNAs were defined as transcripts exhibiting partial or complete overlap with annotated genes on the opposite strand, whereas *trans*-encoded sRNAs were defined as transcripts originating from intergenic regions without overlap with annotated genes. For subsequent downstream analyses, *cis*-encoded sRNAs overlapping tRNA and rRNA genes were removed from further consideration.

The same thresholds applied for gene-level analysis were used to identify differentially expressed sRNAs (padj < 0.05 and |LFC| > 1). Subsequently, differentially expressed *cis*-encoded sRNAs were subjected to an operon filtration step using operon predictions with OperonMapper (Taboada et al. 2018). If a *cis*-encoded sRNA overlapped multiple genes within the same operon, only the gene with the greatest overlap was selected as the representative target and the others were excluded, whereas if the overlapping genes belonged to different operons, none were excluded from further analysis. Pearson correlation coefficients were calculated to assess the relationship between filtered *cis*-encoded sRNAs and target expression levels. To account for multiple hypothesis testing, p-values were adjusted again using the BH procedure for false discovery rate (FDR) control. Only sRNA–target pairs with statistically significant correlation (FDR

< 0.05 and |r| > 0.5) were selected and visualized with heatmaps, displayed in a neck-to-neck orientation with their target sequences (**Supplementary File S4**).

**Supplementary Note S2. Fourier-transform infrared spectroscopy.**

Fourier-transform infrared (FTIR) spectroscopy was performed for chemical characterization of the extracted polymer. Infrared spectra were recorded using a Nicolet 6700 FTIR spectrophotometer (Thermo Fisher Scientific) operated in single-bounce attenuated total reflectance (ATR) mode with a Smart iTR accessory. Measurements were performed using a diamond crystal plate with a 42° angle of incidence. For each spectrum, 32 scans were collected over the range of 600–4000 $cm^{-1}$ at a resolution of 4 $cm^{-1}$. Poly(3-hydroxybutyrate-co-3-hydroxyvalerate) (PHBV; 2% 3-HV; Sigma-Aldrich) was analyzed as a reference polymer. Spectra were cropped to the 1900–700 $cm^{-1}$ region and normalized to a maximum absorbance of 1 within this range.

**Supplementary references**

Ahmad, Shadab, et al. (2025), 'The UniProt Website API: Facilitating Programmatic Access to Protein Knowledge', *Nucleic Acids Research*, 53/W1: W547–53, https://doi.org/10.1093/nar/gkaf394.

Alexa, A., and J. Rahnenführer (2025), 'topGO: Enrichment Analysis for Gene Ontology.', https://doi.org/10.18129/B9.bioc.topGO.

Binns, David, et al. (2009), 'QuickGO: A Web-Based Tool for Gene Ontology Searching', *Bioinformatics*, 25/22: 3045–6, https://doi.org/10.1093/bioinformatics/btp536.

Cantalapiedra, Carlos P., et al. (2021), 'eggNOG-Mapper v2: Functional Annotation, Orthology Assignments, and Domain Prediction at the Metagenomic Scale', *Molecular Biology and Evolution*, 38/12: 5825–9, https://doi.org/10.1093/molbev/msab293.

Lawrence, Michael, et al. (2013), 'Software for Computing and Annotating Genomic Ranges', *PLoS Computational Biology*, 9/8: e1003118, https://doi.org/10.1371/journal.pcbi.1003118.

Ozuna, A., et al. (2020), 'Baerhunter: An *R* Package for the Discovery and Analysis of Expressed Non-Coding Regions in Bacterial RNA-Seq Data', *Bioinformatics*, 36/3: 966–9, https://doi.org/10.1093/bioinformatics/btz643.

Taboada, Blanca, et al. (2018), 'Operon-Mapper: A Web Server for Precise Operon Identification in Bacterial and Archaeal Genomes', *Bioinformatics*, 34/23: 4118–20, https://doi.org/10.1093/bioinformatics/bty496.
